# Surfaceome CRISPR activation screening uncovers ligands regulating tumor sensitivity to NK cell killing

**DOI:** 10.1101/2025.09.17.676710

**Authors:** Ravi K. Dinesh, Xiaotong Wang, Imran A. Mohammad, Pari Gunasekaran, Kostas Stiklioraitis, Jeremy R. Villafuerte, Anirudh Rao, Rogelio A. Hernandez-Lopez, John B. Sunwoo, Le Cong

## Abstract

Natural killer (NK) cell-based immunotherapies represent a promising avenue for cancer treatment due to their ability to eliminate cancer cells independently of antigen presentation and potential for “off-the-shelf” use. However, the molecular determinants governing tumor cell susceptibility to NK cell-mediated cytotoxicity remain incompletely understood. Here we employed CRISPR activation (CRISPRa) screening to systematically identify cancer cell surface regulators of NK cell killing across multiple cancer types. Using a comprehensive surfaceome-focused library, we screened human and murine cancer cell lines co-cultured with NK cells, identifying both known and novel ligands that modulate NK cell cytotoxicity. Our screens revealed established factors including CD43 (encoded by *SPN*), while uncovering previously uncharacterized regulators such as CD44, PDPN, and Siglec-1/CD169. Validation through complementary cDNA overexpression and genetic knockout approaches confirmed that disruption of CD43, CD44, PDPN, and Siglec-1 significantly altered cancer cell susceptibility to NK killing both *in vitro* and in humanized mouse models. Analysis of clinical datasets show that expression of identified factors correlates with patient survival outcomes in an NK-context dependent manner supporting their therapeutic relevance. Most notably, our mechanistic studies demonstrate that CD43-mediated NK cell resistance operates independently of its previously proposed interaction with Siglec-7 on NK cells. Furthermore, we find that targeting CD43 on either NK cells or engineered T cells substantially enhances their cytotoxic activity against leukemia cell lines. These results establish gain-of-function screening as a powerful approach for discovering immunoregulatory surface proteins and identify multiple promising targets for enhancing NK cell-based cancer immunotherapies.

## Introduction

NK cells are a type of innate lymphoid cell (ILC) capable of detecting cellular stress and eliminating cells that are infected or transformed^1^. In contrast to CD8 T cells which kill transformed cells in an antigen-dependent manner via peptide-MHC, NK cell killing is antigen-independent and determined by the balance of activating and inhibiting ligands on the surface of target cells^1^.

To date, cancer immunotherapy research has primarily focused on targeting immune checkpoint interactions that inhibit CD8 T cell killing or engineering T cells using chimeric antigen receptor (CAR) technology^2,3^. NK cells act orthogonally to CD8 T cells by targeting cancer cells that have downregulated surface MHC to evade T cell recognition. Thus, targeting tumor-NK interactions represents a promising alternative or complementary approach to established cancer immunotherapies. Additionally, CAR NK cells offer the promise of “off-the-shelf” manufacturing with decreased risk of cytokine release syndrome (CRS) and graft-versus-host-disease (GVHD) compared to CAR T cells^4,5^. Despite promising clinical trials of interventions focused on NK cells^6–8^, adoption of NK cell therapies in clinical settings is still in its infancy^8,9^. Consequently, studies that elucidate new tumor-NK interactions and pathways offer potential for the discovery of new interventions.

Previous genome-wide loss-of-function screens in cancer cell lines co-cultured with NK cells have identified genes affecting cancer cell sensitivity to NK cell killing^10–15^. However, these approaches were limited to interrogating only endogenously expressed genes. Additionally, a previous study that validated hits from parallel genome-wide and targeted surfaceome screening has demonstrated that surfaceome-focused approaches reduce noise while increasing true positives and decreasing false negatives^16^.

Checkpoint blockade, which largely targets cell surface interactions between immune cells and the tumor microenvironment (TME), remains the most clinically successful immunotherapy^17^, suggesting that uncovering novel interactions driving NK cell dysfunction in the TME might be especially promising as synergistic or complementary modalities^18^.

With this in mind, we sought to identify tumor cell surface regulators of NK cell killing across cancers by utilizing gain-of-function screening methodologies. We employed CRISPR activation (CRISPRa) to screen a library of single guide RNAs (sgRNA) targeting the cancer surfaceome^19^ in multiple cell lines, identifying putative ligands expressed on cancer cells and stroma that modulate sensitivity or resistance to NK cells. We validate the key findings using a complementary cDNA overexpression system and follow with studies demonstrating that knockout of CD43, CD44, PDPN, and Siglec-1/CD169 in cancer cell lines alters their susceptibility to killing by NK cells both *in vitro* and *in vivo* in humanized transgenic mice. For our most robust hit, CD43, we demonstrate that expression of its canonical immune receptor Siglec-7/CD328 on NK cells is dispensable for its ability to promote resistance to NK killing. Noting CD43 expression on primary NK cells and T cells, we further demonstrate that ablation of CD43 on immune cells themselves enhances killing of leukemia cell lines. Our studies reveal novel candidate ligands on cancer cells and tumor stroma that modulate NK cell activity and, importantly, identify potential targets for cancer immunotherapy development.

## Results

### Gain-of-function surfaceome screening identifies known and novel ligands that drive resistance and sensitivity to NK cell cytotoxicity

We were interested in discovering novel ligands expressed across various cancers that could serve as potential targets for NK-based immunotherapy. First, K562-CRISPRa cells were generated with vectors encoding components of the synergistic activation mediator (SAM) system^20,21^ (Fig. S1a) and a clone with stable expression of vector components and high levels of gene activation activity was selected for screening (Fig. S1b). We then created a high complexity customized library of sgRNAs targeting all predicted cancer surfaceome genes (Supplementary Data 1) and transduced them into K562-CRISPRa cells. Library cells were co-cultured with or without primary NK cells for 24 hours with the surviving cells collected, processed for genomic DNA, and sequenced for sgRNA enrichment (Fig. 1a).

**Fig. 1:**
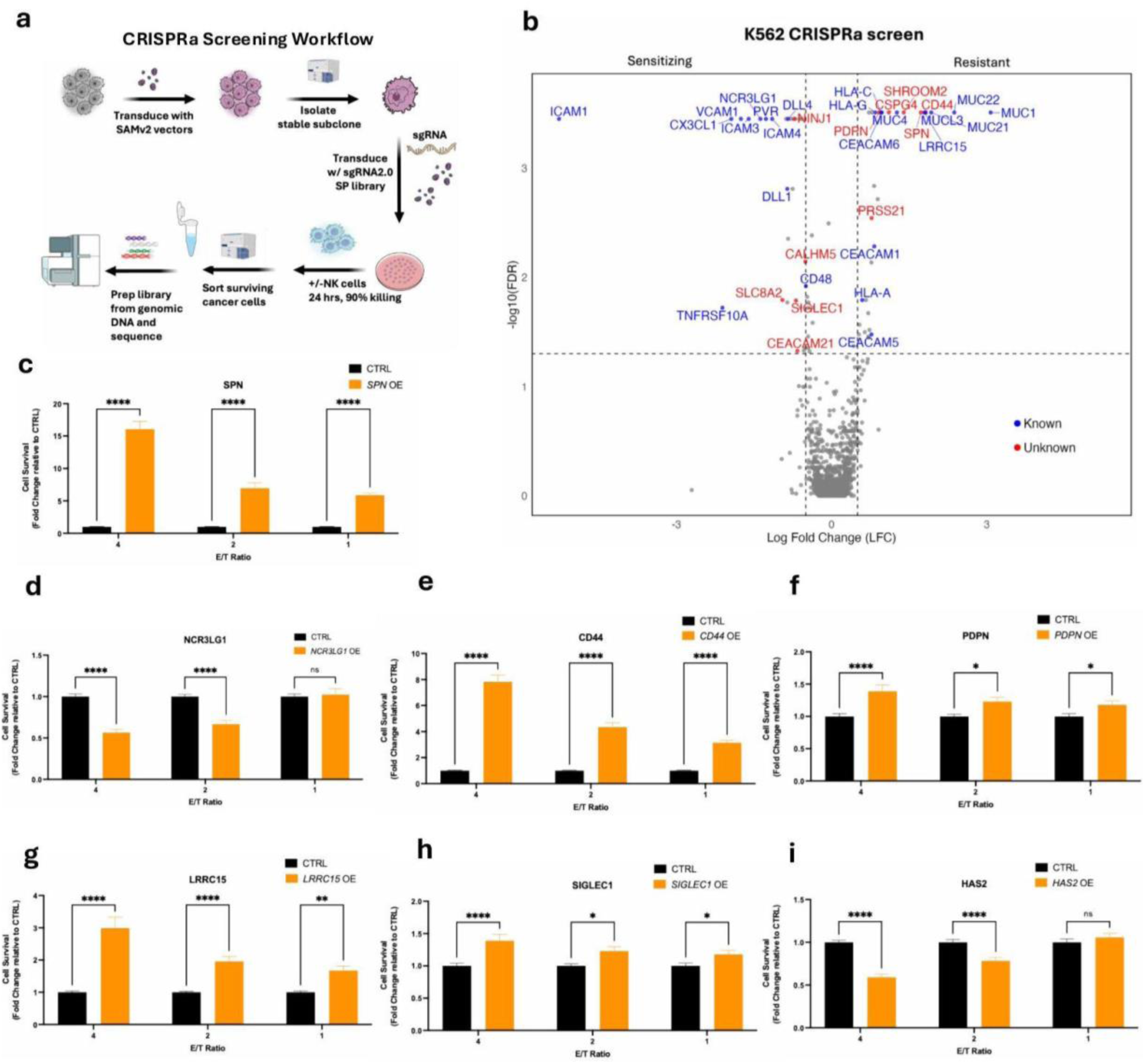
A surfaceome CRISPRa screening identifies putative ligands regulating NK cell cytotoxicity. **a,** Diagram of screening workflow. **b,** Volcano plot of surfaceome CRISPRa screen in K562 CRISPRa cells. Known screen hits appear in blue and novel screen hits in red. **c-i,** NK cell cytotoxicity assays performed using primary NK cells against K562-GFPFluc cells overexpressing the cDNAs SPN (**c**), NCR3LG1 (**d**), CD44 (**e**), PDPN (**f**), LRRC15 (**g**), SIGLEC1 (**h**), and HAS2 (**i**) at different E/T ratios. Data for each ratio is normalized to CTRL. Three to four independent experiments, with 6 technical replicates each. Two-way ANOVA with correction for multiple comparisons. *, p<0.05; **, p<0.01; ***, p <0.001; ****, p<0.0001

Results from our screen in K562 cells demonstrated that our workflow could be used to find both known and unknown sensitizing and resistance factors of NK-mediated killing, such as *NCR3LG1* (encoding B7-H6), mucin family genes, *PVR*, ICAM family genes, *CD244*, *CEACAM6*, and HLA family genes (Fig. 1b). The screen also uncovered a number of novel hits that were not found in previous screens using loss-of-function methodologies^11,13–15,22^ such as the putative sensitizing gene *SIGLEC1* and the resistance genes *CD44*, *SPN*, *PDPN*, and *LRRC15*. As a complementary approach, we also conducted surfaceome CRISPRa screens using a CRISPRa subclone of the human gastric carcinoma cell line HGC-27 (Fig. S1c), which provided confirmation of hits from our K562 screen, such as *NCR3LG1*, *CD48*, *CXC3L1*, *MUC1*, and *LRRC15* as well as uncovering novel genes, such as the hyaluronan synthase *HAS2* (Fig. S2a).

We had hypothesized that the CRISPRa–as opposed to the CRISPRko– approach would have the advantage of discovering genes not expressed endogenously in the cell line tested. To see if this was borne out by our screen results, we used DepMap RNA-Seq data to examine the expression of hits from our two screens in their respective cell lines (Fig. S2b-e). For our K562 screens, ∼61% of our screen hits were expressed at transcripts per million (TPM) < 1 and ∼31% were expressed at near undetectable levels (TPM < 0.1) in the K562 cell line (Fig. S2d).Though not as robust, our HGC-27 screen was also able to uncover genes outside of its endogenous expression context with ∼30% of screen hits expressed at TPM < 1 (Fig. S2e).

As NK cells from human and mouse have divergences in their development and surface receptors^23^, we were interested to see the degree to which our screen hits might be conserved in a murine context as conserved genes might serve as stronger potential therapeutic targets. We conducted screens in the murine lymphoma line YAC-1 as well as the murine melanoma line B16F0, and found similarly strong enrichment for *Cd244*, *Cd48*, Icam, Ceacam, and Mucin family members as well as MHC genes (Fig. S3a,d).

*Siglec1* appeared as a strong sensitizing hit in both screens, suggesting its importance as a putative novel regulator of NK responses (Fig. S3a-b,d). We then compared the strongest overlapping hits across the four screens (Fig. S3c-d) and mapped their expression across TCGA cancers (Fig. S3e). These results further underscored the ability of our approach to uncover genes across cancers and outside the context of the cell lines screened.

We selected seven genes for further validation: the sensitizing genes *NCR3LG1*, *HAS2*, and *SIGLEC1* and the resistance genes *SPN*, *CD44*, *LRRC15*, and *PDPN*. *NCRL3LG1,* a highly established sensitizing ligand for the NK receptor NKp30 (encoded by *NCR3*)^22,24,25^ and *SPN* (which encodes CD43), a recently discovered resistance ligand that has been shown to bind the receptor Siglec-7 on NK cells^26,27^, were selected as positive controls. The other genes were selected for the strength of their overlaps in our screens (Fig. S3d) and/or their potential novelty as ligands for receptors on NK cells. We chose a complementary validation approach using overexpression of cDNAs in a K562 line with a GFP-luciferase expression cassette knocked-in to the *AAVS1* safe harbor locus (hereafter K562-GFPFLuc) (Fig. S4a-b). Results from NK cytotoxicity assays performed with these overexpression lines (Fig. S4e-f) directionally recapitulated the effects seen in our CRISPRa screen, though the relative magnitude was not the same in all cases (Fig 1b-i). Overexpression of *SPN* and *CD44*, in particular, showed much stronger effects using our overexpression system than was seen in CRISPRa screens (Fig 1b-c,e). In total, these results demonstrate the utility of gain-of-function surfaceome screening in identifying novel factors that alter susceptibility to NK cell cytotoxicity.

### Disruption of candidate factors in endogenous contexts recapitulates altered cancer cell susceptibility to NK cell killing in vivo and in vitro

We were next interested in understanding if disruption of endogenous expression of our candidate factors would lead to converse effects on NK cell cytotoxicity. We transduced cancer cell lines with LentiCRISPRV2 vectors encoding Cas9 and either gRNAs targeting our genes of interest or the control *OR10A2* locus. We then sorted the cells to create homogenous negative (candidate factor) and positive (*OR10A2*) populations.

Overexpression of CD43 (encoded by the *SPN* gene) in our validation studies increased resistance to cancer cells to the highest levels seen among all tested candidate factors (Fig. 1c). CD43 is a heavily glycosylated protein whose expression is largely found on cells of hematopoietic origin^28^ and in the context of cancers is found most prominently on leukemias and lymphomas with reports of expression in some colon and salivary gland cancers^29,30^. As K562 cells expressed high levels of CD43 expression on their surface, we tested CD43(+) and CD43(-) K562 cells (Fig. 2a) against primary cells. Strikingly, we saw a close to four-fold increase in killing of the CD43(-) cells at the highest E/T ratio (Fig. 2b). Likewise, when we tested CD43(+) and CD43(-) OCI-AML2 cells (Fig. S5a), we saw a greater than 5-fold effect at the highest E/T ratio (Fig. S5b). Using a humanized NOD-scid-IL2rg null mouse with transgenic expression of IL-15 (NSG-TghuIL15 mice)^31^, we found no difference in tumor growth between CD43(+) and CD43(-) K562 tumors *in vivo* in the absence of NK cells, but adoptive transfer of primary NK cells led to CD43(-) tumors having significantly compromised growth in response to NK cell killing (Fig. 2c).

**Fig. 2:**
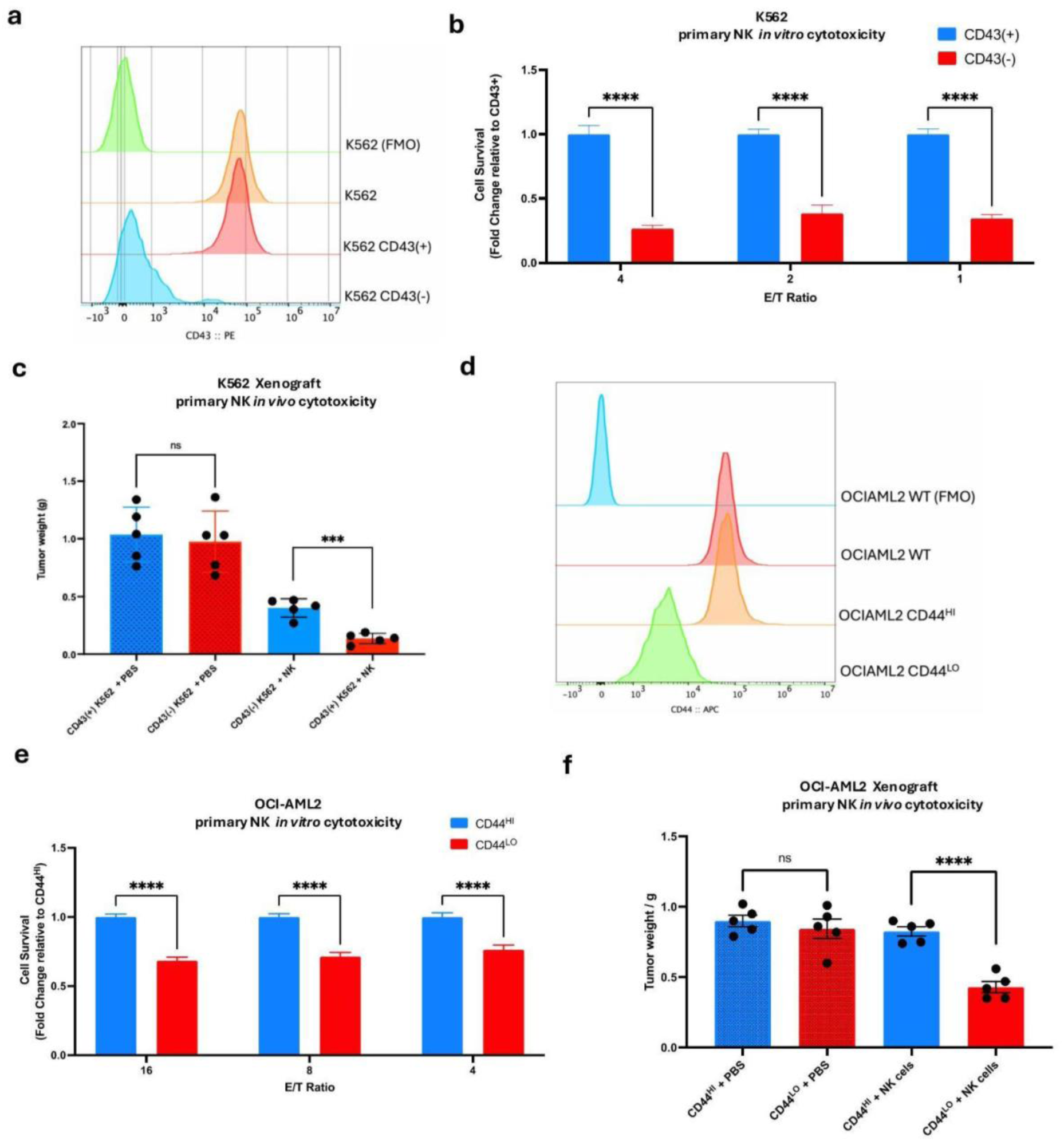
Ablation of CD43 and CD44 increases susceptibility to NK cells *in vivo* and *in vitro*. **a,** Flow cytometry analysis plot of K562 cells and derivatives for surface expression of CD43. **b,** NK cell cytotoxicity assays performed against CD43(+) and CD43(-) K562 cells. Data normalized to CD43(+) for respective E/T ratio. **c,** Day 17 tumor weight from sacrificed NSG-TghuIL15 mice engrafted s.c. with 1 x 10^6^ K562 cells (Day 0) followed by s.c. injections of 3 x 10^6^ NK cells (D3, D6). **d,** Flow cytometry analysis plot of OCI-AML2 cells and derivatives for surface expression of CD44. **e,** NK cell cytotoxicity assays performed against CD44^HI^ and CD44^LO^ OCI-AML2 cells. Data normalized to CD44^HI^ for respective E/T ratio. **f,** Day 19 tumor weight from sacrificed NSG-TghuIL15 mice mice engrafted s.c. with 1 x 10^6^ OCI-AML2 cells (Day 0) followed by s.c. injections of 4 x 10^6^ NK cells (D2, D4). **b,e,** Three independent experiments, 6 technical replicates each. Two-way ANOVA with correction for multiple comparisons. **c,f,** n=5 mice per group, Two-tailed T test with Welch’s correction. *, p<0.05; **, p<0.01; ***, p <0.001; ****, p<0.0001

CD44, like CD43, is a large, heavily glycosylated protein that is widely expressed on immune cells and thought to play a role in immune cell adhesion and signaling^32^. CD44 had the second largest effect on NK cell cytotoxicity in our initial overexpression validation assays (Fig. 1e). As CD44 expression is widespread in the immune system, we initially attempted to ablate CD44 in multiple leukemia lines, including K562, THP1, and RCH-ACV, but found in nearly every case transduction of cells with gRNAs targeting CD44 were slow to recover and showed no evidence of CD44 loss (data not shown). Only OCI-AML2 cells recovered following transduction and featured a hypomorphic phenotype short of a full knockout (Fig. 2d). In cytotoxicity assays, these hypomorphic CD44^LO^ cells demonstrated significantly increased susceptibility to NK cells compared to their CD44^HI^ counterparts (Fig. 2e). This effect was also found *in vivo* where CD44^LO^ cells were similarly compromised in their resistance to NK killing (Fig. 2f).

These data implicate both CD43 and CD44 in resistance to NK cytotoxicity and provide strong evidence for their potential as immunotherapy targets.

### Ligands commonly expressed on tumor stroma alter NK cell susceptibility

Three of the genes validated in our arrayed cDNA expression studies–*SIGLEC1*, *PDPN*, and *LRRC15*–are expressed as much or more frequently on tumor stroma than they are on tumor cells themselves. Both PDPN and LRRC15 are expressed on cancer associated fibroblasts^33–35^ and Siglec-1/CD169 expression is largely confined to tissue resident and tumor infiltrating macrophages^36,37^.

At baseline, MV4-11 cells, a cell line with macrophage-like characteristics, express cell surface CD169 (encoded by *SIGLEC1*) barely measurable above background (Fig. 3a). However, we found that treatment with IFN-α substantially increased levels of CD169 on the cell surface after 48 hrs (Fig. 3a) before returning to being undetectable once IFN-α was removed from culture (data not shown). We transduced MV4-11 cells with LentiCRISPRv2 vectors targeting the *SIGLEC1* or *OR10A2* locus and induced surface Siglec-1 expression with IFN-α before sorting for negative and positive populations, respectively. Following extended culture in IFN-α free media, we assayed both 48hr IFN-α treated and untreated CD169(-) and CD169(+) cells in co-culture with NK cells and found that knockout of *SIGLEC1* lead to increased resistance of MV4-11 cells (Fig. 3b-c). Importantly, pretreatment with IFN-α led to increased resistance by CD169(-) cells demonstrating that expression of CD169 on the cell surface was important for mediating the effect of *SIGLEC1* knockout (Fig. 3b-c). Experiments in NSG-TghuIL15 mice implanted with IFN-α treated MV4-11 cells demonstrated that loss of *SIGLEC1* also increased resistance to NK cytotoxicity *in vivo* (Fig. 3d).

**Fig. 3:**
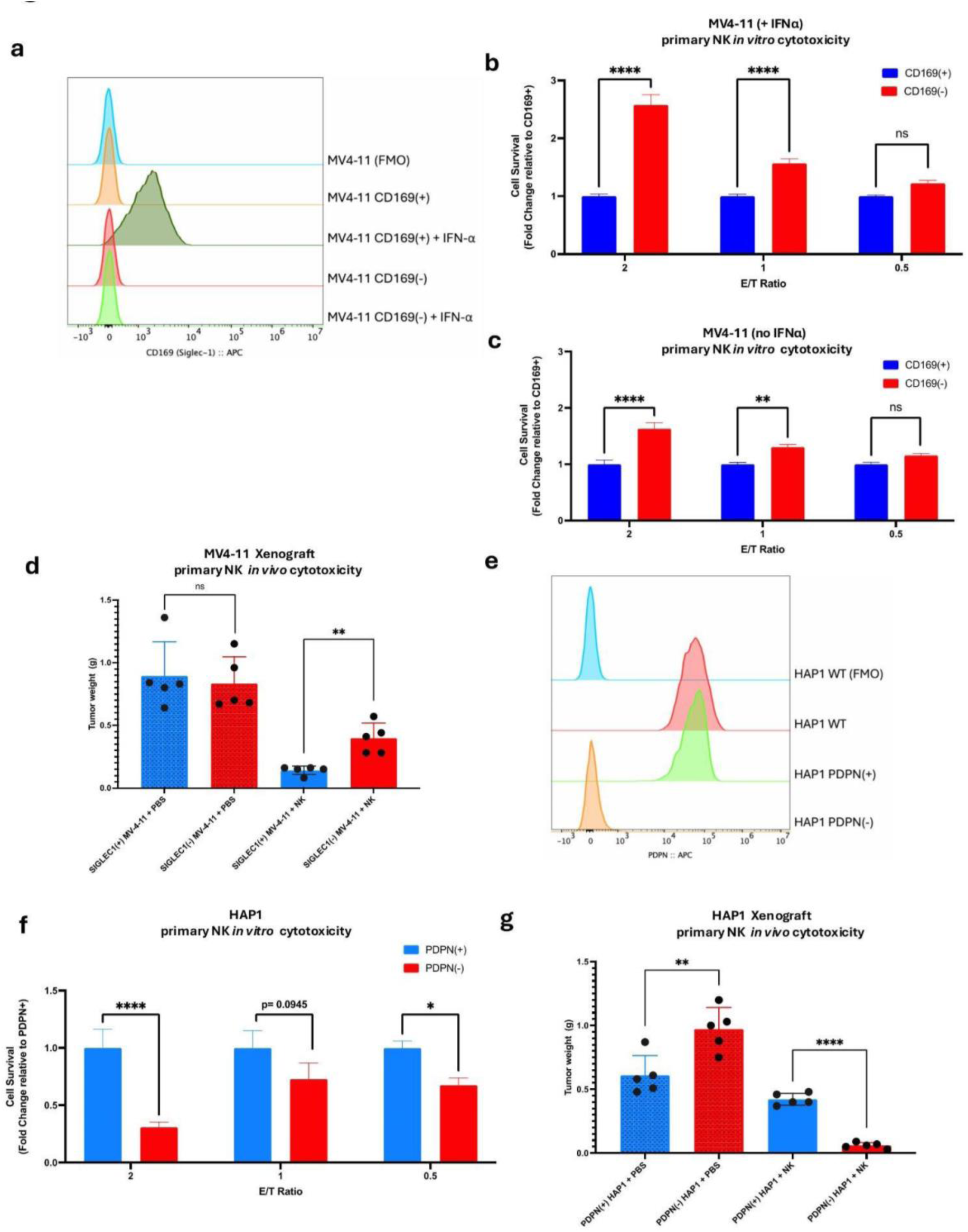
Loss of *SIGLEC1* or *PDPN* alters NK cell responses *in vivo* and *in vitro*. **a,** Flow cytometry analysis plot of MV4-11 cells and derivatives for surface expression of CD169 with and without IFN-α (10 ng/mL). **b,c** NK cell cytotoxicity assays performed against CD169(+) and CD169(-) MV4-11 cells with (**b**) and without (**c**) 48 hr pre-treatment with IFN-α (10 ng/mL). Data normalized to CD169(+) for respective E/T ratio at 24 hr. time point. **d,** Day 34 tumor weight from sacrificed NSG-TghuIL15 mice mice engrafted s.c. with 1 x 10^6^ MV4-11 cells (Day 0) followed by s.c. injections of 2 x 10^6^ NK cells (D3, D6). **e,** Flow cytometry analysis plot of HAP1 cells and derivatives for surface expression of PDPN. **f,** NK cell cytotoxicity assays performed against PDPN(+) and PDPN(-) HAP1 cells. Data normalized to PDPN(+) cells for respective E/T ratio at 24 hr. time point. **g,** Day 31 tumor weight from sacrificed NSG-TghuIL15 mice mice engrafted s.c. with 1 x 10^6^ HAP1 cells (Day 0) followed by s.c. injections of 2 x 10^6^ NK cells (D3, D6). **b-c,f,** Three independent experiments, 6 technical replicates each. Two-way ANOVA with correction for multiple comparisons. **c,f** n=5 mice per group, Two-tailed T test with Welch’s correction. *, p<0.05; **, p<0.01; ***, p <0.001; ****, p<0.0001

We found that the HAP1 cell line expressed high levels of PDPN (Fig. 3e) and was amenable to prolonged culture and genetic manipulation. Loss of PDPN (Fig. 3e) made HAP1 cells significantly more susceptible to NK cell killing *in vitro* (Fig. 3f). When implanted into NSG-TghuIL15 mice *in vivo*, PDPN(-) HAP1 cells grew at a faster rate compared to PDPN(+) cells in mice untreated with NK cells (Fig. 3g). Strikingly, treatment of implanted NSG-TghuIL15 mice with adoptively transferred NK cells strongly reversed this phenotype with PDPN(+) cells showing a minimal effect of NK treatment and PDPN(-) becoming close to eliminated (Fig. 3g). This suggested that despite PDPN-mediated inhibition of tumor growth *in vivo*, its expression is likely selected for in the TME in the context of an immune response to cancer cells.

CD169 and PDPN–unlike CD43 and CD44–are not expressed on NK cells. This allowed us to take advantage of previously developed clinical analysis that examines correlates between tumor gene expression, NK infiltration, and patient survival^38^. We used CIBERSORTx^39^ to classify TCGA cancers into NK rich and NK poor groups (defined as cancer at the top 10% and bottom 90% of NK enrichment, respectively) and examined differences in patient survival when comparing high and low expression of our genes of interest (defined as the top and bottom 50% by expression) (Fig. 4a). Among patients in the NK Rich group, *SIGLEC1* High tumors showed an increased likelihood of survival relative to *SIGLEC1* Low with a Hazard Ratio (HR) of 0.716 (95% CI = 0.574- 0.894) (Fig. 4b-c). This trend was reversed in the NK Poor group which showed decreased survival for *SIGLEC1* High tumors (HR = 1.350, 95% CI = 1.254-1.452) (Fig. 4b-c). Conversely, *PDPN* High tumors showed a statistically significant decrease in survival in both the NK Rich group (HR = 1.753, 95% CI = 1.399-2.196) and NK Poor group (HR 1.606, 95% CI=1.491-1.728) with only a small increase in HR in NK Rich relative to NK poor, indicating that this method could not not find a strong NK-specific effect on decreased survival due to *PDPN* expression (Fig. 4d-e). To offer mechanistic insight into these effects, we then examined if either *SIGLEC1* or *PDPN* was affecting the activation states of NK cells within the tumor. We plotted gene expression from TCGA against the fraction of activated NK cells in tumors as determined by CIBERSORTx. Increasing *SIGLEC1* expression in tumors showed a moderate correlation with increased NK activation (R = .242, p < .001) (Fig. S6a). Conversely, *PDPN* expression was negatively correlated with NK activation (R = -.027, p < .001), though this effect was minimal (Fig. S6b). Taken together, these analyses broadly support the directional effects found for *SIGLEC1* in our *in vitro* and *in vivo* NK cell cytotoxicity experiments.

**Fig. 4:**
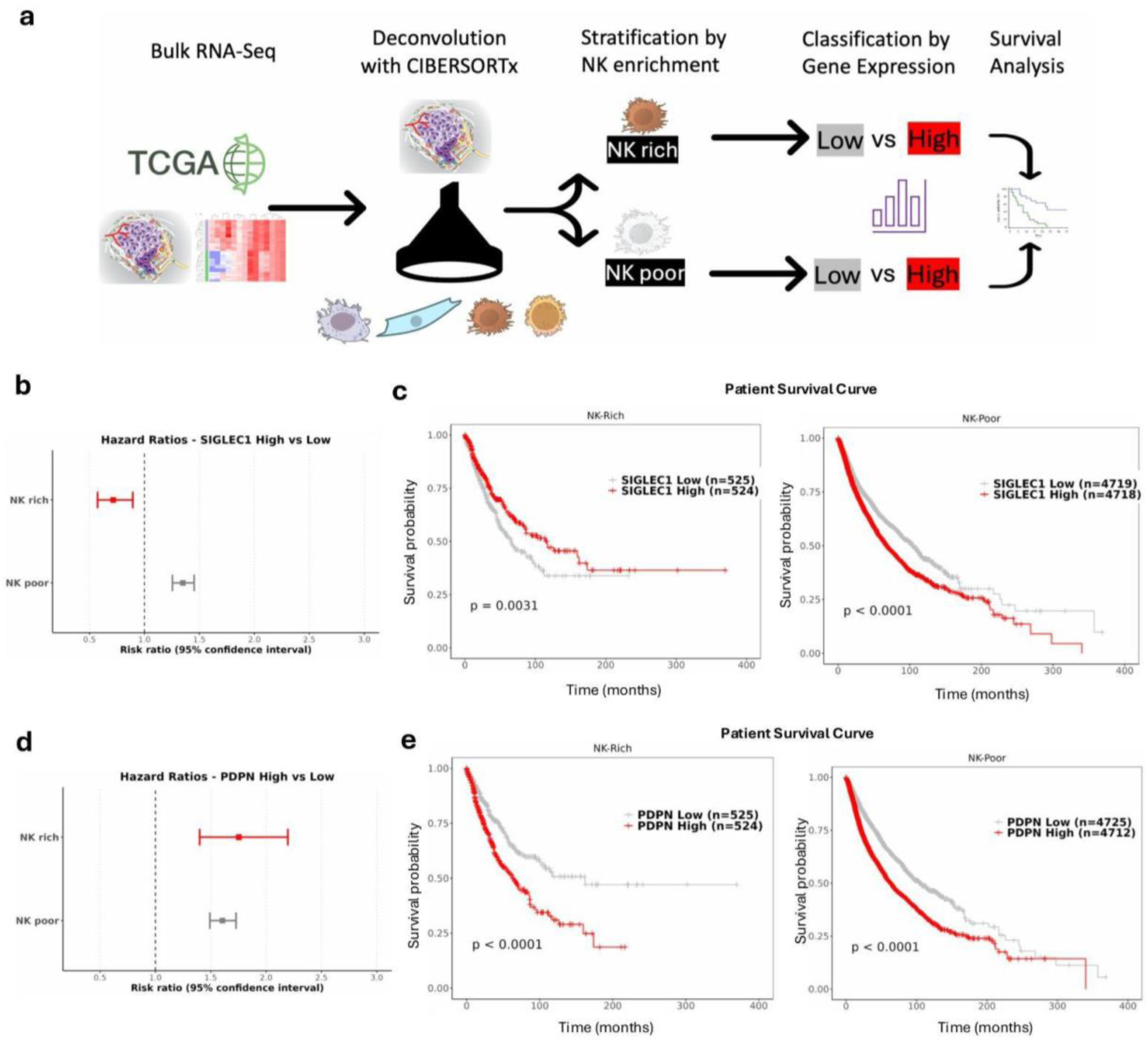
NK-context specific gene expression predicts patient survival. **A,** Diagram of CIBERSORTx and survival analysis of TCGA data. **b,d,** Cox hazard ratios of *SIGLEC1* (**b**) or *PDPN* (**d**) expression stratified by top 10% and bottom 90% NK enrichment in TCGA cancers. **b,d,** Kaplan-Meier plot of *SIGLEC1* (**c**) or *PDPN* (**e**) expression stratified by top 10% and bottom 90% NK enrichment in TCGA cancers.

As CD8 T cells perform a similar cytotoxic function as NK cells as part of the adaptive arm of the immune system, we were also interested to see if similar effects could be found between CD8 Rich and Poor groups. Unlike in the NK rich group, we found no statistically significant difference for *SIGLEC1* in the CD8 Rich group (HR = .903, 95% CI = 0.713-1.144), while the CD8 Poor group (HR = 1.367, 95% CI = 1.271-1.470) showed a decrease in survival for *SIGLEC1* high tumors (Fig. S6c-d). Similar to the NK analysis, both CD8 Rich and CD8 Poor groups showed decreases in survival in the *PDPN* High tumors with CD8 rich (HR = 2.062, 95% CI = 1.616-2.630) showing stronger decreases compared to CD8 Poor (HR = 1.612, 95% CI = 1.498-1.734) (Fig. S6e-f). This data indicates that the effects of *SIGLEC1* are not generalizable to the presence of cytotoxic cells more broadly and are mechanistically acting in a potentially NK-specific manner. For *PDPN* stratification, both NK Rich and CD8 Rich tumors have small-to-moderate increases in hazard ratios relative to their Poor counterparts (Fig. 4d-e, S6e-f), suggesting that *PDPN* expression has broader effects on immune cell populations that cumulatively affect tumor responses and patient survival.

We conclude from these experiments and analyses of clinical data that both Siglec-1/CD169 and PDPN potentially have important roles in dictating immune responses in the TME and further validate gain-of-function screening as a method to find immunoregulatory ligands expressed on tumor stroma.

### Siglec-7 on NK cells is dispensable for CD43-mediated repression of NK killing

Among all candidates validated, overexpression (Fig. 1c) or knockout (Fig. 2b-c, S5b) of *SPN* had the largest effects on NK cell killing of target cells. Thus, we were interested in better understanding the mechanisms by which *SPN* mediated its effects. Two recent reports^26,27^ have shown that CD43 binds Siglec-7 and claim that this interaction is responsible for SPN’s repressive effects on NK cells.

Curiously, previous studies had reported that, unlike primary NK cells, the NK-92 cell line–which is being examined in numerous clinical trials as a cell immunotherapy treatment^8^--had either undetectable levels of Siglec-7^40^ or very low levels in the absence of extended culture with horse-serum containing media^41^. We cultured our NK- 92 and NK-92MI lines in the absence of horse serum and confirmed with two different anti-CD328/Siglec-7 antibodies that these cell lines expressed very low levels of Siglec- 7 relative to primary NK cells with NK-92MI cells having close to undetectable levels (Fig. 5a, S7a-c). We also examined SIGLEC7 transcript expression in NK-92, NK-92MI, and primary NK subsets from peripheral blood using published RNA-Seq data which confirmed our staining results (Fig. 5b, S7d).

**Fig. 5:**
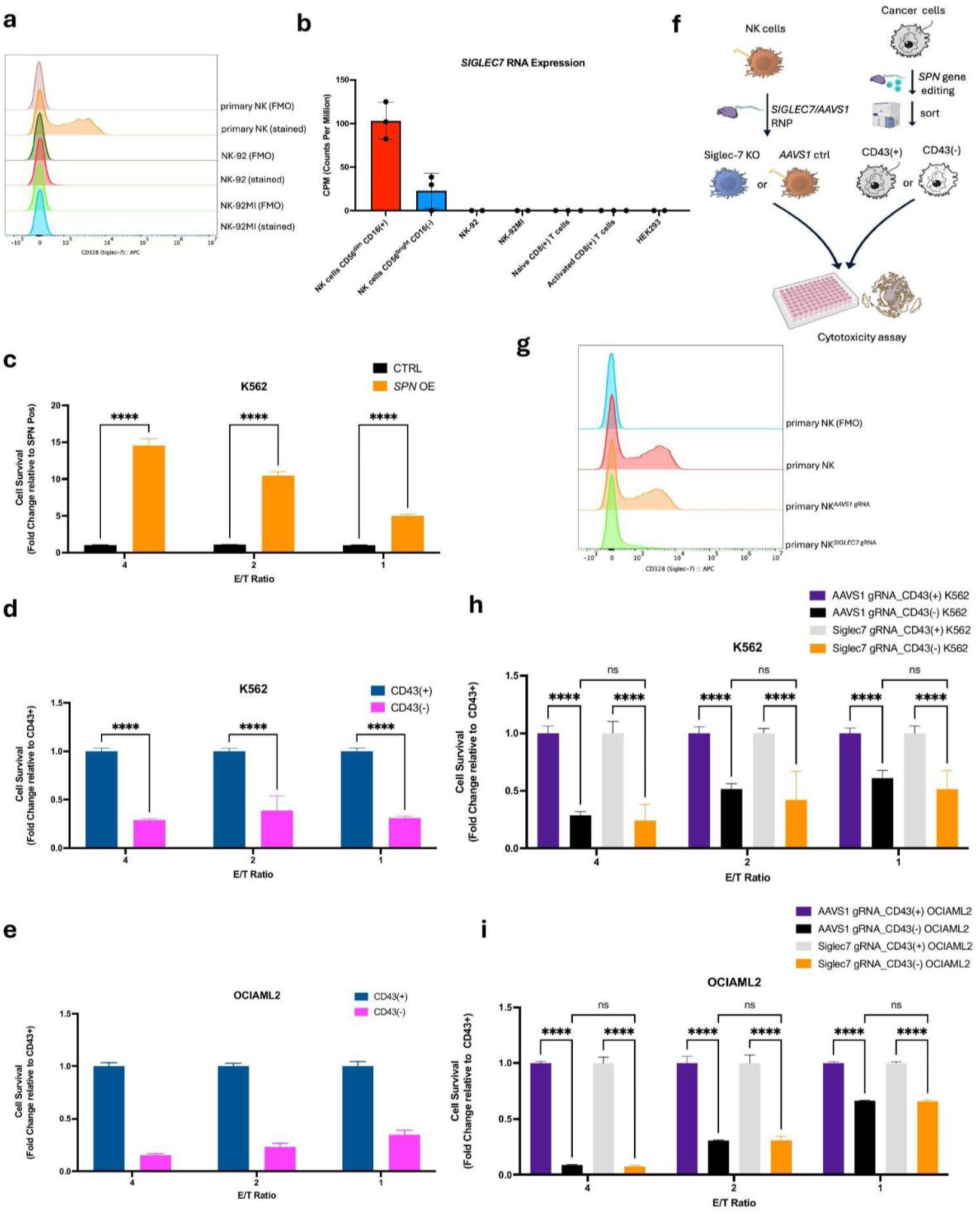
Siglec-7 is dispensable for CD43-mediated repression of NK cell cytotoxicity. **a,** Flow cytometry analysis of CD328 expression in primary NK cells, NK-92, and NK-92MI. **b,** Normalized expression of *SIGLEC7* transcripts from published RNA-Seq data. Primary NK cell subsets were derived from PBMC. **c,** NK-92MI cytotoxicity assays performed against K562-GFPFluc cells overexpressing SPN or control cells. Data normalized to CTRL for respective E/T ratio at 24 hr. time point. **d-e,** NK-92MI cytotoxicity assays performed against CD43(+) and CD43(-) K562 (**d**) or OCI-AML2 (**e**) cells. Data normalized to CD43(+) cells for respective E/T ratio at 24 hr. time point. **f,** Workflow of gene edited primary NK cell cytotoxicity experiments. **g,** Representative flow cytometry analysis of CD328 expression in RNP-targeted primary NK cells four days after nucleofection. **h-i,** NK cell cytotoxicity assays performed against CD43(+) and CD43(-) K562 (**h**) or OCI-AML2 (**i**) cells using *AAVS1* or *SIGLEC7* targeted primary NK cells. Data normalized to CD43(+) cells for respective E/T ratio at 24 hr. time point. **b**, n=3 for primary NK groups, and n=2 for cell lines. **c,** Five independent experiments, 6 technical replicates each. **d-e,** Three independent experiments, 6 technical replicates each. **h-i,** Two independent experiments, 6 replicates each. Two-way ANOVA with correction for multiple comparisons. *, p<0.05; **, p<0.01; ***, p <0.001; ****, p<0.0001

We hypothesized that these Siglec-7^Lo^ NK-92MI cells would be less responsive to differential levels of CD43 on the surface of cancer cells. Strikingly, cytotoxicity assays performed with Siglec-7^Lo^ NK-92MI cells against K562 SPN cDNA overexpressing cells demonstrated a strong positive effect of CD43 on NK cell killing (Fig. 5c). This effect was further corroborated by results comparing stimulation by CD43(-) versus CD43(+) K562 cells (Fig. 5d) or by CD43(-) versus CD43(+) OCI-AML2 cells (Fig. 5e). When comparing the effect of cancer cell SPN ablation on NK cytotoxicity as measured by fold change in survival between CD43(-) cells relative to CD43(+) cells, we found no significant differences between different E/T ratios using primary NK cells or NK-92MI with the exception of the 1:1 ratio for OCI-AML2 (Fig. S7e-f). Here, the difference between primary NK and NK-92MI was in the opposite direction expected if CD43 mediated its repressive effects through Siglec-7 (Fig. S7f). To rule out the possibility that Siglec-7 was upregulated on NK-92MI cells in the presence of cancer cells, we co-cultured NK-92MI cells with K562 cells for 24 hr and measured surface Siglec-7 levels but found no changes that would explain the lack of expected effect (Fig. S3g-h).

Since we could not rule out the possibility that CD43 acts differentially on primary NK cells and NK-92MI or through multiple pathways, we then performed cytotoxicity experiments using primary NK cells where *SIGLEC7* was targeted by CRISPR ribonucleoprotein (RNP) (Fig. 5f). Over multiple experiments, we were able to see ablation of Siglec-7 expression at 70-99% when compared to cells transfected with a control AAVS1 RNP (Fig. 5g, S7i). In NK cytotoxicity assays, Siglec-7 ablated primary NK cells showed no differences in their responsiveness to differential CD43 expression on either K562 (Fig. 5h) or OCI-AML2 (Fig. 5i) cells when compared to control AAVS1- targeted NK cells.

From these various lines of evidence, we conclude that CD43 binding to Siglec-7 on NK cells is not the primary mechanism through which its immune repressive effects are mediated.

### Targeting CD43 on primary NK cells improves cytotoxic responses against cancer cells

CD43 is highly expressed on immune cells, including NK cells and T cells (Fig. S8b,d). As we had ruled out Siglec-7 as a dominant factor mediating CD43 repression, we hypothesized that targeting SPN on immune cells themselves could drive improved cancer cell killing.

We utilized MEM-59, a CD43-binding monoclonal antibody that had previously been validated in blockade experiments in cancer cells^26^ (Fig. S8a). Remarkably, blockade of CD43 on NK cells followed by cytotoxicity assays against the leukemia lines K562, Nalm6, OCI-AML2, or SUP-B15 demonstrated significantly improved NK killing (Fig. 6a-e). As treatment with NK cells or CAR NK cells has emerged as a cell therapy treatment modality^42^, we were interested to see if genetic ablation of CD43 could improve primary NK killing of leukemia cells. We used CRISPR RNP to ablate CD43 on primary NK cells (Fig. 6f, S8b) and found that–similar to blockade–genetic targeting of *SPN* significantly improved NK killing against leukemia lines when compared to the control *AAVS1*-targeted cells (Fig. 6g-j).

**Fig. 6:**
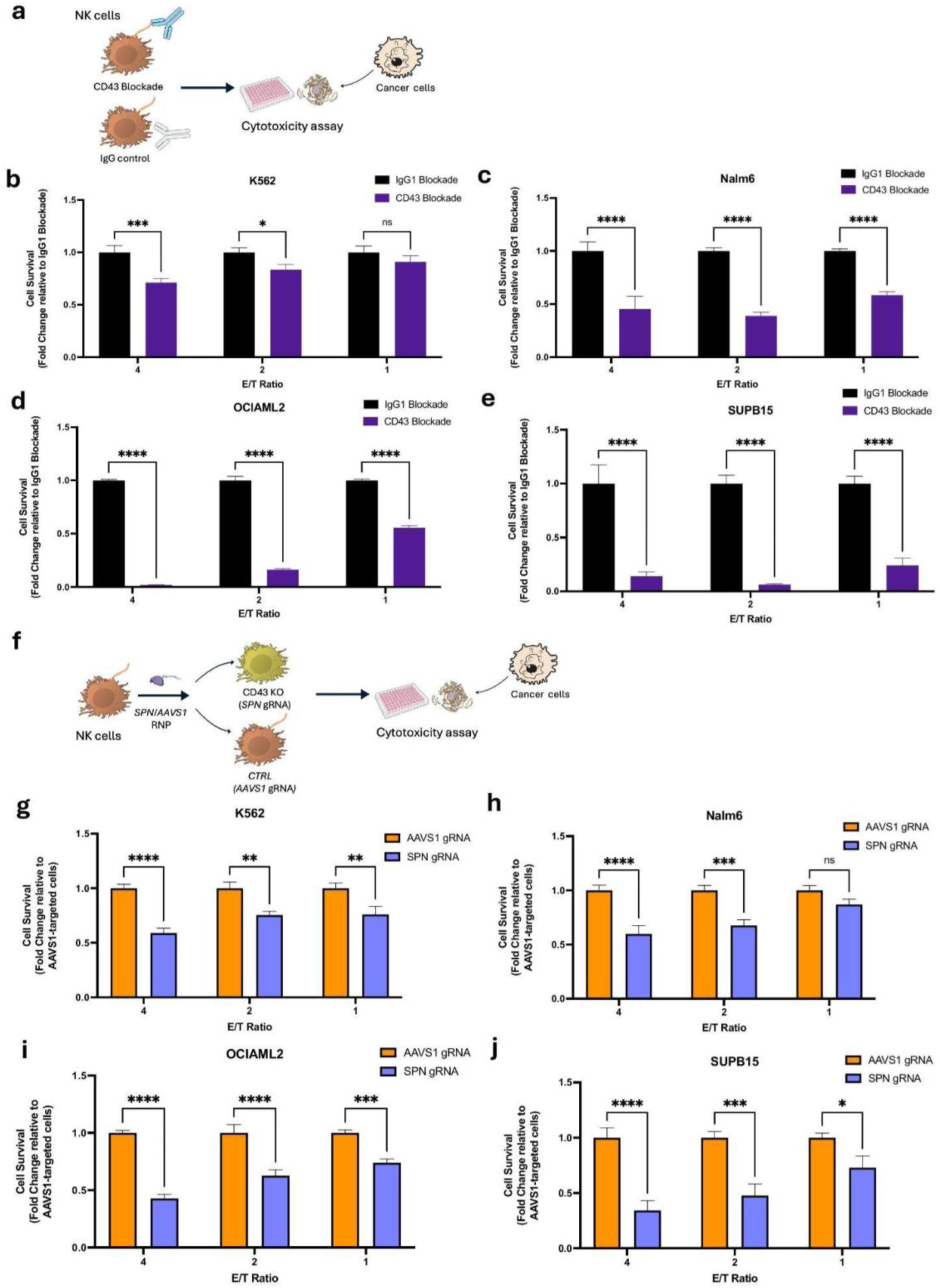
Targeting CD43 on NK cells improves killing of leukemia cells. **a,** Workflow of cytotoxicity experiments utilizing blockaded NK cells. **b-e,** Cytotoxicity assays performed with blockaded primary NK cells against K562 (**b**), Nalm6 (**c**), OCI-AML2 (**d**) or SUPB15 (**e**) cells. Data normalized to control mouse IgG1-blockaded cells for respective E/T ratio at 24 hr. time point. **f,** Workflow of cytotoxicity assays performed with gene edited primary NK cells. **g-j,** Cytotoxicity assays performed with RNP-targeted primary NK cells against K562 (**g**), Nalm6 (**h**), OCI-AML2 (**i**) or SUP-B15 (**j**) cells. Data normalized to control AAVS1-targeted cells for respective E/T ratio at 24 hr. time point. **b-d,** n=4, **e,** n=2, **g,i-j,** n=2 **h,** n=3 independent experiments, 3-6 replicates each. Two-way ANOVA with correction for multiple comparisons. *, p<0.05; **, p<0.01; ***, p <0.001; ****, p<0.0001

Like NK cells, T cells also have substantive CD43 expression on their cell surface (Fig. S8d). As modification of CAR T cells through genetic perturbation or exogenous cDNA expression has shown promise in many pre-clinical studies^43–50^, we hypothesized that targeting CD43 on CD8 T cells would improve their activity against leukemia cells similarly to NK cells. We transduced CD8 T cells with a CD19 targeting CAR and assayed them against CD19(+) lines Nalm6 and SUP-B15 (Fig. 7a, S8c). Like primary NK cells, CD19-targeted CAR T cells showed improved killing upon CD43 blockade (Fig. 7a-b). We next used CRISPR RNP to target *SPN* on CD19-targeted CAR T cells before sorting *SPN* and control *AAVS1*-targeted cells for CD43(-) and CD43(+) populations respectively (Fig. 7c, S8d). In cytotoxicity assays, CD43(-) CD19-targeted CAR T cells showed significant and substantial improvements in cytotoxicity over their CD43(+) counterparts (Fig. 7d-e).

**Fig. 7:**
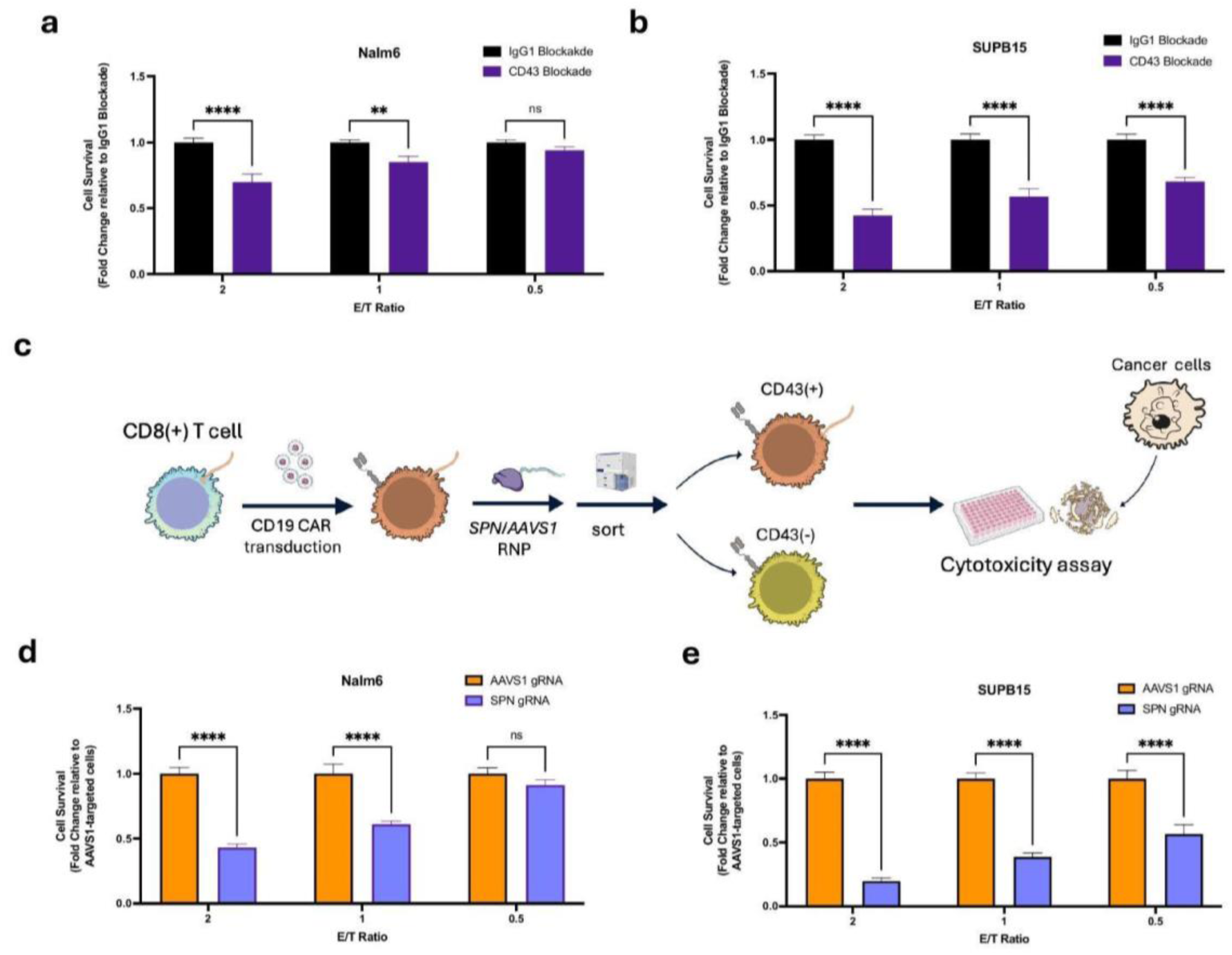
Targeting CD43 on CD19-targeting CAR T cells increases killing against CD19(+) cells. **a-b,** Cytotoxicity assays performed with blockaded CD19-targeted CAR T cells against Nalm6 (**a**) or SUPB15 (**b**) cells. Data normalized to control mouse IgG1-blockaded cells for respective E/T ratio at 24 hr. time point. **c,** Workflow of cytotoxicity experiments using gene edited CAR T cells. **d-e,** Cytotoxicity assays performed with RNP-targeted and sorted CD19-targeted CAR T cells against Nalm6 (**d**) or SUP-B15 (**e**) cells. Data normalized to control AAVS1-targeted CD43(+) cells for respective E/T ratio at 24 hr. time point. **a-b,** n=5, **d-e,** n=2 independent experiments, 3 replicates each. Two-way ANOVA with correction for multiple comparisons. *, p<0.05; **, p<0.01; ***, p <0.001; ****, p<0.0001

Taken together, we demonstrate that targeting CD43 on either primary NK cells or CD8 T cells can improve cytotoxic activity. This suggests that targeting CD43 through blockade or in the context of cell therapy could show promise as a therapeutic against leukemia.

## Discussion

In the work presented we apply gain-of-function methodology to identify putative novel surface proteins that act to alter sensitivity of cancer cells to NK cell cytotoxicity across cancer cell types and in tumor stroma. Using a comprehensive validation approach utilizing cDNA overexpression, knockouts in cell lines with endogenous expression, and *in vivo* experiments in NSG-TghuIL15 mice we are able to demonstrate that the genes *CD44*, *SPN*, *PDPN*, and *SIGLEC1* have critical roles in cancer cell susceptibility to NK cell killing.

Previous screens probing cancer cell determinants of NK cell susceptibility used CRISPR knockout approaches with genome-wide libraries^11,13–15,22,51^. Based on our past work^52^ and work from other groups^16,53,54^, we reasoned that a gain-of-function CRISPRa approach with a library targeting surface proteins would reduce the noise associated with genome-wide screens and increase the potential for finding ligands outside of those expressed endogenously in the cancer line tested. These assumptions were validated by our screens, which identified genes outside of the endogenous context of the cancer cell lines screened, including in stromal and immune cells associated with the TME.

While this manuscript was in preparation, Dufva *et. al.*^55^ published genome-wide screens in blood cancer lines co-cultured with NK cells that included genome-wide CRISPRa screens. Their results did not point to significant enrichment of genes such as *SIGLEC1* or *PDPN* that we identified in our surfaceome screens^55^. Though we cannot rule out lab-specific variations in protocol, we believe this is best explained by the previously noted potential for domain-specific sub-pooled libraries to outperform genome-wide libraries in finding true positives^16^.

Of the genes we pursued for extensive validation, CD169/Siglec-1 has largely been studied in the context of viral infections, where it mediates pathogen uptake, pathogen recognition, and cross-presentation to dendritic cells^37^. Its role remains understudied in cancer, but work has demonstrated that ratio of CD169(+)/CD68(+) macrophages predicts improved clinical outcomes in gliomas and breast cancer^56,57^ and its increased expression serves as a biomarker of positive outcomes in malignant melanoma^56^. We provide evidence here that Siglec-1 is not just a marker of a beneficial macrophage population, but that it acts functionally to improve NK cell cytotoxicity against tumors and that this correlates strongly with improved clinical outcomes across cancers. As such, we believe therapies that seek to recruit populations of CD169(+) stromal cells or increase expression of CD169(+) in the tumor stroma, as has been shown with TLR9 agonists in viral contexts^56^, will serve to improve cancer immunotherapy.

Overexpression or ablation of *SPN*/CD43 among genes tested in our validation studies had the largest impact on resistance to NK cell killing. A number of ligands and coreceptors have been implicated as interactors with CD43, including E-selectin^58,59^, CD169^60^, ICAM1^61^, and the cytoskeletal-interacting ERM (Ezrin/Radixin/Moesin) proteins^62,63^. These interactions were largely thought to play a role in mediating CD43- dependent cell trafficking with its role in lymphocyte activation determined by intracellular mechanisms^64^.

More recently, independent work from two groups has demonstrated that genetic ablation or antibody blockade of CD43 on target cells spurs improved NK cell killing^26,27^. These studies further identified a biochemical interaction between Siglec-7 on natural killer cells and CD43 on cancer cells that was determined in part by the sialylation status of CD43 and suggested this interaction was the primary determinant of CD43- mediated resistance to NK cell cytotoxicity^63,65^. Subsequent publications on Siglec-7 or CD43 have taken the functionality of this interaction as a given^55,66–68^. However, we demonstrate here that lack of detectable expression of Siglec-7 in an NK cell line line does not alter CD43-mediated resistance. More importantly, we show that genetic disruption of Siglec-7 in primary NK cells similarly has no effect on responsiveness to CD43. One study reported a functional effect of the CD43/Siglec-7 interaction by simultaneously overexpressing SPN on K562 cells and Siglec-7 on NK-92 cells^27^. In contrast, these NK-92 *SIGLEC7* overexpression cells showed no differential cytotoxicity against *SPN*-knockdown K562 cells or K562 cells expressing endogenous levels of SPN^27^. As such, the data presented here cannot rule out the possibility that tandem supraphysiological expression of CD43 and Siglec-7 may lead to some functional effects on NK or T cell cytotoxicity, but this is unlikely to be the primary mechanism through which CD43 mediates it effects.

We further demonstrated that blockade or genetic ablation of CD43 on either NK cells or T cells could improve killing against leukemia cell lines. Previous work has suggested that targeting either CD43 or Siglec-7 would allow for improved immune control of CD43(+) tumors^26,27^. However, our results suggest that any beneficial effects of targeting either molecule would be independent of the other, though it’s possible they could act additively. Additionally, our data indicate that therapies targeting CD43 on leukemias and lymphomas, especially, could function synergistically by acting on both cancer cells and immune cells simultaneously.

In sum, the work presented here serves as a demonstration of the utility of focused gain-of-function screening to identify surface proteins on cancer cells and tumor stroma that can alter NK cell cytotoxicity. More importantly, we show through extensive validation that a number of these factors serve as promising targets for improved NK cell-based immunotherapies.

## Materials and Methods

### Plasmids and vector construction

LentiCRISPRv2 (Addgene #52961), lentiCas9-Blast (Addgene #52962), and dCas9VP64_GFP (Addgene #61422) were gifts from Feng Zhang. AAVS1 SA-2A-puro-pA (Addgene #22075) was a gift from Rudolf Jaenisch. pXPR_502 (Addgene #96923) was a gift from John Doench and David Root. pSLCAR-CD19-BBz (Addgene #135992) was a gift from Scott McComb. pD649-HAsp-SIGLEC1(Trunc1)-COMP5AP-AviTag- 9xHis (Addgene #157610) was a gift from Chris Garcia. espCas9(1.1)-2A-GFP Gln tRNA plasmids containing gRNAs targeting AAVS1^69^ were gifts from David Schatz.

pLV-EF1a-dCas9VP64_2A_EGFP_2A_BSD, pLV-EFS-PCP_p65_HSF1_2A_mcherry_HygR, pLV-U6_sgRNA_pp7-EFS-Puro_2A_BFP, pLV- 2Apuro_2A_BFP, AAVS1-BSD-EF1a_Fluc2_2A_GFP, and pLV-EF1a- Fluc2_2A_GFP_2A_BSD, pLV-puro_2A_BFP, and all cDNA containing plasmids were generated by Gibson Cloning using the NEB Builder HiFi DNA Mastermix (New England Biolabs E2621L). pLV-EF1a-dCas9VP64_2A_EGFP_2A_BSD was generated by inserting a 2A-BSD fragment into dCas9VP64_GFP cut with BsrgI and EcoRI. pLV-EFS-PCP_p65_HSF1_2A_mcherry_HygR was generated by inserting EFS, PP7_p65_HSF1 (from pXPR_502), 2A_mcherry, and 2A_HygR fragments into EF1a- EGFP-2A-ZeoR^52^ cut with PacI/EcoRI. pLV-U6_sgRNA_pp7-EFS-Puro_2A_BFP was generated by inserting EFS, Puro, and 2A_mtagBFP fragments into pXPR_502 cut with XhoI/MluI. pLV-2A_puro_2A_BFP was generated by inserting 2A_puro and 2A_BSD fragments into EF1a-EGFP-2A-ZeoR cut with MluI/BamHI. All cDNA constructs were generated by amplifying fragments from commercially purchased cDNAs (Supplementary Data 3) and inserted into pLV-2A_puro_2A_BFP cut with BamHI. AAVS1-BSD-EF1a_Fluc2_2A_GFP was generated by inserting P2A_BSD, SV40polyA- EF1a, Fluc2 (Firefly luciferase 2), and 2A_EGFP fragments into AAVS1 SA-2A-puro-pA cut with XhoI/PmeI. pLV-EF1a-Fluc2_2A_GFP_2A_BSD was used to create Fluc2 expressing cancer cell lines (with the exception of K562) for cytotoxic assays and was generated by inserting FLuc2_2A_GFP and 2A_BSD into pLV-puro_2A_BFP.

CAR sequences were built by fusing the CD19 scFv to the hinge region of the human CD8α chain and transmembrane domains, and cytoplasmic regions of the human 4-1BB, and CD3z signaling domains. The anti-CD19 scFV (FMC63) sequence was amplified from pSLCAR-CD19-BBz. The CAR sequences were cloned into a modified pRRLSIN vector powered by human phosphoglycerate kinase (hPGK) promoter and flanked upstream by N-terminal CD8a signaling peptide (MALPVTALLLPLALLLHAARP) and membrane targeting myc-tag (EQKLISEEDL), and downstream by an (SASGSG)-P2A-mScarlet3 sequence.

gRNA vectors were generated by ordering sequences as pairs of complementary oligos either from Thermo Fisher or Integrated DNA Technologies, annealing in 1X T4 DNA Ligase Buffer (New England Biolabs #B0202S) using a thermocycler ramp down protocol after heating to 95C, and Golden Gate cloned into LentiCRISPRv2 with NEBridge Golden Gate Assembly Kit (BsmBI-v2) (New England Biolabs# E1601). A list of gRNA targets and sequences can be found in Supplementary Data 4.

### Cell Culture

293FT cells (#R7007) were purchased from Thermo Fisher Scientific. HGC-27 cells (# 94042256) were purchased from Sigma Aldrich. B16F0 (#CRL-6322), YAC-1 (#TIB-160), and NK-92MI (#CRL-2408) cell lines were purchased from ATCC. K562 cells were a gift from Michael Cleary (Stanford University). OCI-AML2 cells were a gift from Rizwan Romee (Dana-Farber Cancer Institute). HAP1 cells were a gift from Jan Carette (Stanford University). MV4-11 cells were a gift from Kathleen Sakamoto (Stanford University). SUPB15 cells were a gift from Kara Davis (Stanford University). Nalm6-GFP cells were a gift from Crystal Mackall (Stanford University).

293FT and B16F0 cells were grown in DMEM (Corning #10-101-CV) supplemented with 1% Pen-Strep (Gibco 15-140-122), 1% NEAA (Cyclone #SH30238.01) and 10% FBS (BenchMark) at 37°C. K562 cells were grown in RPMI (Corning #10-104-CV) supplemented with 1% Pen-Strep, 1% NEAA, and 10% FBS at 37°C. HGC-27 cells were grown Eagle’s Minimum Essential Medium (Quality Biological #112-016-101CS) supplemented with 1% Pen-Strep, 1% NEAA, and 10% FBS at 37°C. All other cell lines were grown in NK media: RPMI supplemented with 1% Pen-Strep, 1% NEAA, 1 mM sodium pyruvate (Thermo Scientific #11-360-070), 10 mM HEPES (Cytiva #SH30237.01), and 55 nM 2-mercaptoethanol (Fisher #21985023). NK-92 and NK-92MI cells were grown in NK media and supplemented with 100 IU/mL of IL-2 (NIH NCI).

### Lentiviral production and transduction

Lentiviruses were generated by transfecting 293FT cells at 80% confluence using JetOptimus (Polyplus #117-01) and a DNA ratio of 2:2:1 transfer plasmid:pCMV- dR8.91:pCMV-VSVG according to manufacturer’s protocol. Media was replaced after 4 hours and viral supernatant harvested at 48 hours. Viral supernatant was passed through a 0.45 uM filter to clear debris and was either used immediately or stored at -80 C. Surfaceome libraries were pooled together from multiple plates and concentrated 10X using Lentivirus Precipitation Solution (Altstem #VC100) according to manufacturer’s protocol and stored in -80 C. Cells were transduced in 6, 12, or 24 well plates with viral supernatant using polybrene (Millipore Sigma #TR-1003) at a final concentration of 8 ug/mL and spinfected at 1000 x g for 45 min at 32 C. For surfaceome screening, concentrated virus was titrated on 1 x 10^6^ cells in a 6 well plate at a final volume of 3 mL using the aforementioned transduction reagents and spinfection conditions. The percentage of cells that were BFP+ was determined by FACS at 48 hours and was used to calculate the MOI. For the screen, viral supernatant corresponding to MOI 0.5 was used to transduce cells.

### Lentiviral production and transduction of human T cells

Pantropic VSV-G pseudotyped lentivirus was produced by transfecting Lenti-X 293T cells (Clontech #11131D) with a pRRLSIN transgene expression vector and the viral packaging plasmids pCMVdR8.91 and pMD2.G using PEI (Polysciences, Inc. #24765100). Primary T cells were thawed and after 24 hours in culture, were stimulated with Human T-Activator CD3/CD28 Dynabeads (Life Technologies #11131D) at a 2:1 cell:bead ratio. After 48 hours, viral supernatant was harvested and added to primary T cells. T cells were exposed to the virus for 24 hr after which virus containing supernatant was replaced with fresh media. At day 5 after T cell stimulation, the Dynabeads were removed. T cells were stained and sorted on an Aria Fusion cell sorter (BD Biosciences) on the basis of mScarlet3 expression to obtain homogenous receptor expression levels. Untransduced T cells were used to set up the sorting gates but were not sorted as the test articles. T cells were expanded for at least 9 days and cultured at 0.5 million/mL.

### Surfaceome library construction

For the human surfaceome library, 8 gRNAs per gene for every human cancer surface protein gene as delineated by Hu et. al.^19^ was generated using the CRISPick tool from the Broad Institute. For the mouse surfaceome library, genes from the human surfaceome library were converted to their mouse homologs using the babelgene R package and a library of 8 gRNAs/gene was generated using the CRISPick tool. For each library an oligo pool was ordered from Genscript and cloned into the pLV- U6_sgRNA_pp7-EFS-Puro_2A_BFP vector using Golden Gate cloning with BsmBI as described previously^52^. Briefly, isopropanol precipitation of the library was performed before transformation, and precipitated library DNA was transformed into MegaX DH10BTM T1R ElectrocompTM Cells (Thermo Fisher #C640003) with electroporation for amplification. After Maxi prep, NGS was performed for library quality control. Sequences for gRNA libraries can be found in Supplementary Data 1.

### Generation of CRISPRa screening lines

K562, HGC27, YAC-1, and B16F0 CRISPRa lines were generated by first transducing cells with pLV-EF1a-dCas9VP64_2A_EGFP_BSD and selecting with blasticidin (Invivogen #ant-bl-05) (20 ug/mL for HGC-27 and B16F0, 10 ug/mL for K562 and HGC27) for 3-7 days. The cells were then transduced with pLV-EF1a- PCPp65HSF1_2A_mcherry_HygR and selected in hygromycin (Invivogen #ant-hg-5) (125 ug/mL for K652, 500 ug/mL for A375, 1000 ug/mL for YAC-1, and 1 mg/mL for B16F0). Bulk cells were single-cell sorted by FACS for GFP+/mcherry+ double positive populations into 96 well plates. Colonies were picked 10-21 days later and expanded into 24 well plates. Clones with highly stable GFP and mcherry expression after 3-4 weeks in culture (as determined by flow cytometry) that also maintained high levels of CRISPR activation were selected for screening.

### Primary NK cell isolation and culture

Peripheral blood mononuclear cells (PBMCs) were were isolated from LRS chambers obtained from the Stanford Blood Center using Ficoll-Paque (Cytiva #17- 5442-02) and SepMate columns (StemCell Technologies #85450) according to manufacturer’s protocol. Isolated PBMCs were resuspended in PBMC freezing media (90% FBS, 10% DMSO) and stored in liquid nitrogen. Primary NK cells were isolated from thawed PBMCs using the Easy Sep NK cell Isolation Kit (StemCell Technologies #17955) according to manufacturer’s protocol. Isolated NK cells were reconstituted at a density of 1-1.5 x 10^6^ cells/mL of CTS NK-Xpander Medium (Thermo Scientific #A5019001) supplemented with 5% Immune Cell Serum Replacement (Thermo Scientific #A2596101) and containing NK Cell Activation/Expansion beads (Miltenyi Biotec #130-094-483) for six days with additional media added on Day 3. On day 6, activation/expansion beads were removed with an EasySep magnet (Stem Cell #18000) and allowed to expand further in NK media. After isolation NK cells were grown in 1000 IU IL-2 and 20 ng/mL IL-15 (NIH NCI BRB Preclinical Repository).

### Primary human T cell isolation and culture

Primary CD8+ T cells were isolated from the blood of anonymous donors by negative selection (STEMCELL Technologies #17953). T cells were cryopreserved in Cellbanker 1 (Amsbio #11910). T cells were cultured in human T cell medium (HTCM) consisting of X-VIVO 15 (Lonza #04-418Q), 5% Human AB serum, and 10 mM neutralized N-acetyl L-Cysteine (SigmaAldrich #A9165) supplemented with 30 units/mL IL-2 (NCI BRB Preclinical Repository) for all experiments.

### CRISPRa screens in cancer cells co-cultured with NK cells

CRISPRa K562, HGC-27, YAC-1, and B16F0 clonal lines were spinfected with either human (K562, HGC-27) or mouse (YAC-1, B16F0) surface protein library virus at an MOI of 0.5 in 6 well plates at a total volume of 3 mL of media containing 8 ug/mL polybrene. Following spinfection, cells were allowed to rest for 48 hours before puromycin (Invivogen #ant-pr) (1.5 ug/mL for K562, 2.0 ug/mL for HGC-27 and B16F0, and 0.5 ug/mL for YAC-1) was added to the media. Cells in selection were grown out for 7-10 days in 15 cm dishes and the efficacy of selection was determined by monitoring BFP+% by flow cytometry. Library cells that were at least 98% positive were determined to be ready for screening.

One day prior to initiating screening, a fraction of library cells was subjected to a killing assay to determine the optimal E:T ratio for the screen. Adherent cells (HGC-27 and B16F0) were plated at 0.5 x 10^6^ cells/well 24 hours prior to the assay and suspension cells (K562 and YAC-1) were plated at 1 x 10^6^ on the day of the assay.

Differing ratios of primary NK cells were added to the cancer cells at a final volume of 1 mL in NK media and incubated overnight. At 24 hours, adherent cells were detached with 100 uL of TrypLE (Thermo Scientific #12604013) and quenched with 400 uL of media. A 150 uL aliquot of cells was taken from each well and a similar volume from each E/T ratio was run on a flow cytometer to determine the total number of GFP+/mcherry+ events (surviving cells). This number for each E/T ratio was divided by the total number of GFP+/mcherry+ events in conditions where cancer cells were untreated with NK cells to determine the percentage of surviving cells. The E/T ratio that lead to ∼10% of cells surviving was used for the full-scale screen.

A day prior to screening, a total of 30 x 10^6^ adherent library cells were plated on 15 cm dishes at 3 x 10^6^ cells/plate for both NK and no NK conditions. Suspension library cells were plated at 5 x 10^6^ cells/plate on the day of the screen. To the NK condition, primary NK cells were added, and for both conditions media was added to a total volume of 28 mL of NK media with IL-2 and IL-15 for 24 hours. The following day surviving cancer cells from both NK and no NK conditions were collected and sorted by FACS for GFP+/mcherry+ cells. The cells were pelleted at 300 x g for 5 min, the supernatant removed, and stored in -80C for further processing. Two independent biological replicates were performed for each screen and screen condition. Results of screens can be found in Supplementary Data 2.

### Genomic DNA isolation, gRNA amplification, sequencing, and analysis

Quantification of differential gRNA abundance was carried out as described previously^52^. Briefly, genomic DNA was isolated using Quick-DNA Midiprep Plus Kit (Zymo Research #D4075). gRNA sequences were amplified from genomic DNA using the NEBNExt High-Fidelity 2X PCR Master Mix (NEB #M0541) with 50 uL reaction volumes and 2.5 ug of DNA per reaction and mixed staggered primers (Forward Primer: 5’ AATGATACGGCGACCACCGAGATCTACACTCTTTCCCTACACGACGCTCTTCCGATCT[Stagger, 0-7nt]TTGTGGAAAGGACGAAACACC-3’ Reverse Primer: 5’- CAAGCAGAAGACGGCATACGAGAT [8nt- barcode]GTGACTGGAGTTCAGACGTGTGCTCTTCCGATCTTCTACTATTCTTTCCCC TGCACTGT-3’). PCR reactions were pooled and column purified using Monarch PCR and DNA Cleanup Kit (NEB #T1030S). A fraction of the eluate was run on a gel, the band corresponding to the library amplicon excised, and then column purified. Amplified products were quantified using the Qubit 1X dsDNA HS Assay (Thermo Scientific #Q33230), normalized by concentration and sequenced on an Illumina NextSeq using 500/550 v2.5 kits at 1000X coverage per gRNA. Sequencing data was deconvoluted with the bcl2fastq function (illumina) and MAGeCK^70^ was used to produce ranks based on robust rank aggregation (RRA) scores comparing the NK and no NK conditions. Hits were determined as genes showing a false discovery rate (FDR) < 0.05 and a log fold change (LFC) > 0.5.

### Analysis of Screening Data

For overlap analysis of screen hits across cell lines, significantly enriched or depleted genes (|log₂FC| > 0.25, FDR < 0.05) were compiled from four independent screens (K562, HGC27, YAC1, B16F0). Genes shared in two or more conditions were identified using binary presence matrices. UpSet plots were generated using the ComplexUpset R package (v1.3.3) to visualize shared and unique hits across screens. Venn diagrams were drawn using the VennDiagram package (v1.7.3) to depict overlap patterns among gene sets.

Gene expression data were obtained from the Cancer Dependency Map (DepMap) database using the DepMap R package (v1.22.0). Transcripts per million (TPM) expression values were retrieved using the depmap_TPM() function, and cell line metadata were accessed via the depmap_metadata() function. Gene expression of screen hits (defined as |LFC| > 0.5, FDR > 0.05) were analyzed for their respective cell lines: K562 (DepMap ID: ACH-000551) and HGC-27 (DepMap ID: ACH-000847) a gastric cancer cell line. For each cell line, TPM expression values were extracted for all genes of interest. Expression levels were categorized into five groups: Very High (≥10 TPM), High (5-10 TPM), Moderate (1-5 TPM), Low (0.1-1 TPM), and Very Low (<0.1 TPM). All analyses were performed in R using the following packages: depmap (DepMap data access), dplyr (data manipulation), ggplot2 (visualization), tidyr (data reshaping), readr (file I/O), and tibble (data structure utilities).

For analysis of expression across The Cancer Genome Atlas (TCGA) database, genes of interest were identified as those that were hits (defined as |LFC| > 0.25, FDR > 0.05) in at least two of the four screens conducted. RNA-seq expression data were obtained from the TCGA database via the Genomic Data Commons (GDC) using the TCGAbiolinks R package (v2.37.1). Data were queried for "Transcriptome Profiling" with data type "Gene Expression Quantification" using the "STAR - Counts" workflow, targeting both primary tumor samples and solid tissue normal samples across all available cancer types. Expression data for the GOIs was extracted from individual TCGA TSV files (rna_seq.augmented_star_gene_counts.tsv) to extract TPM (Transcripts Per Million), FPKM (Fragments Per Kilobase Million). Only tumor samples with non-missing TPM values were retained for analysis. Expression values were log2- transformed using log2(TPM + 1) to handle zero values and normalize the distribution.

For each gene-cancer type combination, median log2(TPM + 1) values were calculated to provide robust central estimates of expression. A gene × cancer type matrix was constructed, and cancer types with data available for fewer than 50% of analyzed genes were excluded to ensure data completeness. Z-score normalization was applied across cancer types for each gene to enable comparison of relative expression patterns while maintaining gene-specific expression signatures. The normalization was calculated as (x - μ) / σ, where x is the median expression for a gene in a specific cancer type, μ is the mean expression across all cancer types for that gene, and σ is the standard deviation. Heatmaps were generated with the ggplot2 R package. Genes were ordered by functional category (NK cell sensitizing genes followed by NK cell resistant genes) with alphabetical sorting within each category. Cancer types were clustered using hierarchical clustering with Euclidean distance and complete linkage.

Analysis performed in R used the following packages: TCGAbiolinks (TCGA data download and access), readr (file I/O), dplyr (data manipulation), tidyr (data reshaping), ggplot2 (visualization), pheatmap (publication-quality heatmaps), tibble (data structure utilities), stringr (string manipulation), RColorBrewer (color palettes), and viridis (perceptually uniform color scales).

### CIBERSORTx and Survival Analysis

Analyses were based on an approach reported by Oh *et al.*^38^ using R software with the survival, survminer, dplyr, and ggplot2 packages. Gene expression and clinical data concerning 10,524 primary tumors within the Cancer Genome Atlas (TCGA) was downloaded from the UCSC Xena Database^71^. The CIBERSORTx web-application was used to estimate the abundance of NK cells in each cancer sample by providing bulk TCGA gene expression data, and the LM22 matrix file that defines gene expression profiles of 22 immune cell types^39^. CIBERSORTx was run with 500 permutations, without quantile normalization, and in absolute-mode.

For survival analysis, patients were stratified into immune cell-rich and immune cell-poor groups based on total NK cell (sum of activated and resting NK cells) or CD8 T cell abundance. The top 10% of enrichment for a given immune cell was stratified as “Rich” and the bottom 90% as “Poor”. Within each immune stratification group, patients were further divided by gene expression levels using median split (top 50% defined as “High” and bottom 50% defined as “Low”). Survival analysis was performed using the Kaplan-Meier method with log-rank tests to compare survival distributions between gene expression groups. Cox proportional hazards regression was used to calculate hazard ratios and 95% confidence intervals, comparing high vs low gene expression within each immune cell stratum. Survival times were converted from days to months by dividing by 30.42.

Analysis was performed in R using the following packages: dplyr (data manipulation and filtering), tibble (enhanced data frame operations), ggplot2 (data visualization and plotting), survival (survival analysis functions including Cox proportional hazards regression and Kaplan-Meier estimation), ggpubr (publication-ready plot formatting), tidyr (data reshaping and cleaning), and survminer (enhanced survival plot visualization and statistical testing).

### Correlation analysis of gene expression and Immune fractions

Correlation analyses between gene expression and immune cell infiltration were visualized using the packages ggplot2 (v3.4.0), corrplot (v0.95), dplyr (v1.1.0), tidyr (v1.3.0),scales (v1.2.1), and RColorBrewer (v1.1-3). Input data consisted of pre- computed Spearman correlation coefficients between target genes (SIGLEC1, PDPN) versus immune cell fractions (NK_Activated) derived from CIBERSORTx deconvolution results. Gene expression data (FPKM values) from TCGA RNA-seq data downloaded from UCSC Xena were merged with immune cell fraction estimates for approximately 10,524 tumor samples. Correlation heatmaps were generated using the corrplot package with gene expression (FPKM) on the x-axis and immune cell fractions on the y-axis. Due to the wide dynamic range of immune cell fractions, y-axes were log10- transformed and zero or negative immune cell values were replaced with 1×10⁻⁶ to enable log transformation. Linear regression lines with 95% confidence intervals were fitted using geom_smooth() with method="lm".

### Generation of GFP/FLuc2 lines and cDNA expressing derivatives

K562_GFPFLuc cells were created by resuspending 3 x 10^6^ K562 cells in 0.3 mL of RPMI media along with 15 ug of AAVS1-BSD-EF1a_Fluc2_2A_GFP and 7.5 ug each of two espCas9(1.1)-2A-GFP Gln tRNA plasmids targeting AAVS1. The mixture was added to 0.4 cm cuvettes (Bio-Rad #1652088) and electroporated at 875 V/cm^2^, 500 μF, and infinite resistance in a Gene Pulser Xcell Electroporation device^72^. The cells were immediately transferred to 20% FBS containing pre-warmed media and allowed to recover. Five days after transfection, blasticidin (10 ug/mL) was added to the culture to select cells with stable integration of FLuc2/GFP expression cassette. After an additional two weeks in culture, cells with stable integration of the cassette and high expression of GFP were sorted by FACS. cDNA containing cell lines were created by transduction of K562_GFPFLuc cells with pLV-2A_puro_2A_BFP or pLV- 2A_puro_2A_BFP containing genes of interest. Two days later, 2.0 ug/mL puromycin was added to the cells for 5 days to select cells that were successfully transduced. Cells were thereafter maintained in 1.0 ug/mL of puromycin.

OCI-AML2, MV4-11, HAP1, and SUPB15 GFP/Fluc lines were generated by transducing wild type cells with pLV-EF1a-Fluc2_2A_GFP_2A_BSD. After two days, cells were selected with blasticidin (20 ug/mL for OCI-AML2 and SUPB15, and 10 ug/mL for MV4-11 and HAP1). The cells were maintained in blasticidin unless being used in assays.

### Cytotoxicity assays

Adherent cancer cells were plated in either 96 well clear plates or white bottomed tissue culture plates (Grenier Bio-One #655083) at a density of 10 x 10^4^ cells/well one day prior to the assay. Suspension cells were similarly plated at 10 x 10^4^ cells/well on the day of the assay. Primary NK cells, NK-92MI cells, or CAR T cells were added at different effector-to-target (E:T) ratios for each condition and a set of control wells was left without NK cells. For assays with NK cells or NK-92MI cells, wells were filled to 200 uL total of NK media with 1000 IU/mL IL-2 and 20 ng/mL IL-15. For assays utilizing CAR T cells, wells were filled to 200 uL with human T cell medium with 30 IU/mL IL-2. Clear bottomed plates were placed in an IncuCyte live cell analysis system kept inside a tissue culture incubator with readings of fluorescence measured as Total Green Object Integrated Intensity taken every 2 hours. White bottomed plates were returned to a regular TC incubator. At 24 hours, white bottomed plates were removed from the incubator and D-luciferin (Syd Labs #MB000102-R70170) diluted in 1X PBS was added to wells to a final concentration of 1.5 mg/mL and read using the BioTek synergy H4 hybrid microplate reader in endpoint mode . K562 cell lines and their derivatives were analyzed in the IncuCyte, while all other lines were analyzed using a luciferase read- out. Readings at the 24 hour time point were used to calculate differences between empty vector/cDNA or KO/WT groups. Fraction of cells surviving was calculated as: (Signal at X E:T Ratio)/(Signal at 0 E:T ratio) for each condition. To calculate the relative fold change, the fraction of cells surviving for both control/test groups was then divided by the values of empty vector (cDNA overexpression assays) or WT/Pos/HI (endogenous gene ablation assays).

For blockade experiments, NK cells or T cells were incubated in 10 ug/mL of anti-SPN (MEM-59)^65^ (Novus Biologicals NBP2-62227) or mouse IgG1 isotype control (R&D Systems #MAB002) antibody in 1X PBS with 2% FBS or culture media at a concentration of 1 x 10^6^ cells/mL for 20 min at RT. Cells were washed twice with 10X the incubation volume and repelleted in complete media with IL-2 and IL-15 before being used in cytotoxicity assays.

### Generation of CRISPR KO lines

K562, OCI-AML2, MV4-11, or HAP-1 cells expressing GFP and Fluc2 were transduced with LentiCRISPRv2 vectors containing Cas9 and 2-3 gRNAs targeting either a gene of interest (GOI) or the control OR10A2. Two days after transduction, puromycin (2.0 ug/mL for K562 and OCI-AML2, 1.0 ug/mL for MV4-11 and HAP-1) was added to the media and cells were selected for 10 days. Two weeks after transduction, cells were checked by flow cytometry for successful KO of the GOI, pooled together, then sorted by FACS for the KO/Negative phenotype (cells pooled from wells containing gRNAs targeting GOI) or the WT/Positive phenotype (cells pooled from well containing gRNAs targeting OR10A2). Sorted cells were allowed to recover for a week and kept in blastidicin thereafter to maintain GFP/Fluc2 expression before use in assays. Antibodies used for sorting and analysis experiments can be found in Supplementary Data 5.

### In vivo NK cell cytotoxicity assays in NSG-IL15 mice

NSG-TghuIL15 mice (#030890) were acquired from The Jackson Laboratory. All cancer cells were engrafted subcutaneously (s.c.) followed by s.c. delivery of Day 10-14 post-isolation primary NK cells. For experiments using K562 cells, 1 x 10^6^ CD43(+) or CD43(-) K562 cells were engrafted on D0 followed by injections of 3 x 10^6^ NK cells (D3, D6) and tumor measurements on Day 17. For experiments using OCI-AML2 cells, 1 x 10^6^ CD44^HI^ or CD44^LO^ OCI-AML2 cells were engrafted on D0 followed by injections of 4 x 10^6^ NK cells (D2, D4) and tumor measurements on Day 19. For experiments using MV4-11 cells, 1 x 10^6^ CD169(+) or CD169(-) MV4-11 cells were pre-treated with IFN-α (10 ng/mL) (PeproTech #300-02AA) for two days in vitro and then engrafted on D0 followed by injections of 2 x 10^6^ NK cells (D3, D6) and tumor measurements on Day 34. For experiments using HAP1 cells, 1 x 10^6^ PDPN(+) or PDPN(-) HAP1 cells were engrafted on D0 followed by injections of 2 x 10^6^ NK cells (D3, D46) and tumor measurements on Day 34.

### Analysis of *SIGLEC7* expression in RNA-Seq data sets

Raw RNA-seq datasets were downloaded from the NCBI Sequence Read Archive (SRA) using the SRA Toolkit (sratoolkit/2.11.0). The final analysis included 18 samples across seven distinct cell types: primary NK CD56bright CD16(-) cells (n=3; SRR9604267, SRR9604277, SRR9604287), primary NK CD56dim CD16(+) cells (n=3; SRR9604268, SRR9604278, SRR9604288), NK-92 cells (n=2; SRR24900026, SRR24900027), NK-92MI cells (n=2; SRR15898395, SRR15898396), HEK293 cells (n=2; SRR11309003, SRR11309004), naïve CD8 T cells (n=3; SRR17782846, SRR17782880, SRR17782914), and activated CD8 T cells (n=3; SRR17782873, SRR17782907, SRR17782941).

Quality control assessment of raw sequencing reads was performed using FastQC (fastqc/0.11.9) to evaluate read quality scores, GC content, adapter contamination, and other sequence quality metrics. Adapter trimming and quality filtering were performed using Trimmomatic (trimmomatic/0.39) with the following parameters: ILLUMINACLIP with a seed mismatch tolerance of 2, palindrome clip threshold of 30, simple clip threshold of 10, LEADING and TRAILING quality thresholds of 3, SLIDINGWINDOW of 4:15, and MINLEN of 36 to remove reads shorter than 36 base pairs. Trimmed reads were aligned to the human reference genome (GRCh38/hg38) using STAR aligner (star/2.7.9a) with default parameters optimized for paired-end reads. STAR was configured to generate coordinate-sorted BAM files with the --outSAMtype BAM SortedByCoordinate option. The reference genome and gene annotation files (Homo_sapiens.GRCh38.109.gtf) were obtained from Ensembl release 109.

Read counts per gene were quantified using featureCounts from the Subread package (subread/2.0.1) with the following settings: paired-end mode (-p), feature type set to exon (-t exon), gene identifier set to gene_id (-g gene_id), and 8 threads (-T 8) for computational efficiency. Only reads mapping to exonic regions were counted, and genes with fewer than 10 total counts across all samples were filtered out prior to analysis. For visualization purposes, Counts Per Million (CPM) values were calculated by dividing each gene’s raw count by the total library size (in millions) for each sample: CPM = (raw_count × 10^6) / total_library_size. The resulting CPM values were log2- transformed after adding a pseudocount of 1 (log2(CPM + 1)) and then z-score normalized across genes for heatmap generation. Differential expression analysis was performed using DESeq2 (version 1.34.0) in R, which employs a negative binomial generalized linear model and applies variance stabilizing transformations with built-in normalization for library size and composition effects. Genes with adjusted p-values (Benjamini-Hochberg correction) < 0.05 were considered significantly differentially expressed.

Gene expression heatmaps were generated using the pheatmap package in R, displaying z-score normalized expression values across cell types. Genes were manually ordered by functional classification (inhibitory receptors followed by activating receptors) without hierarchical clustering to maintain biological grouping. The analysis focused on a curated panel of 19 immune genes based on their known roles in NK cell and T cell function.

### Generation of primary NK cell and T cell knockouts

gRNAs were ordered from Synthego as EZ sgRNA kits with modified scaffold for *AAVS1* (ACCCCACAGUGGGGCCACUA), *SPN* (AACCCAGAUGAGAACUCACG), and *SIGLEC7* (AGUUCCGUGACCGUGCAAGA) and reconstituted in TE buffer. RNP was generated by combining Alt-R S.p. Cas9 Nuclease V3 (Integrated DNA Technologies) with gRNA at a 1:1 molar ratio and incubated at 37C for 15 min as described previously^73^. Primary NK cells were isolated as described above and cultured in CTS NK-Xpander Medium with 5% Immune Cell Serum Replacement and 100 IU IL-2. Day 6 post-isolation primary NK cells were collected and pelleted in microcentrifuge tubes. Supernatant was removed and NK cells were reconstituted in 40 uL of P3 buffer from the P3 Primary Cell 4D-Nucleofector Kit (Lonza #V4XP-3032) per 10^6^ cells. Twenty uL of NK cells in P3 buffer was added to 2 uL of RNP mix for each reaction, mixed, and pipetted into one well of a 16 well Nucloevette strip. Cells were electroporated with the 4D-Nucleofector using program DN-100 before quickly being returned to 0.5 mL of NK X-pander media with 5% Immune Cell Serum Replacement containing 100 IU IL-2 in 12 well plates. The next day another 0.5 mL of media was added and NK cells were rested for 3 days. Four days after nucleofection, primary NK cells were expanded and checked for successful ablation of the gene of interest. Primary NK cells were then used on 6-8 days post nucleofection (12-14 days post isolation) for downstream assays and analysis.

One day after transduction with CAR-containing vectors, T cells were spun down and reconstituted in 20 uL of P3 Buffer per 10^6^ cells. RNP was generated by mixing 1.56 uL of 100 uM sgRNA (AAVS1 or SPN) and 1.25 uL of 62 uM of Alt-R S.p. Cas9 Nuclease V3 (Integrated DNA Technologies). Twenty uL of the cell mixture was mixed with RNP and electroporated in a 16 well Nucloevette strip on the 4D-Nucleofector using program EO-115. Cells were returned to 1 mL of media in a 12 well plate. Three days later cells were sorted for CD43(+) (AAVS1 gRNA electroporated) or CD43(-) (SPN gRNA electroporated) populations that were also mScarlet3+ (expressing CAR) on an Aria Fusion cell sorter (BD Biosciences). Cells were allowed to rest for an additional six days before use in cytotoxicity assays.

## Author Contributions

R.K.D., L.C., I.A.M., and J.B.S. conceived the study. R.K.D., L.C., X.W., P.G., K.S., J.R.V., R.A.H.-L., and I.A.M. designed experiments. R.K.D., P.G., K.S., X.W., and J.R.V. designed and cloned constructs. R.K.D., P.G., X.W., and K.S. generated cell lines and performed *in vitro* NK cell cytotoxicity assays and associated experiments. R.K.D., X.W. and I.A.M. performed CRISPRa screens. X.W. performed *in vivo* experiments. J.R.V. generated CAR T cells and gene-edited CAR T cells. R.K.D. and J.R.V. performed CAR T cell cytotoxicity assays. R.K.D., P.G., K.S., X.W., J.R.V., and I.A.M. analyzed experimental data. A.R. developed a python script for analysis of Incucyte data. K.S., R.K.D., and X.W. performed bioinformatics analysis. L.C., J.B.S., and R.A.H.-L. supervised experiments. R.K.D. wrote the manuscript with input from the other authors.

## Acknowledgements

This work was supported by the National Institutes of Health grants 1R35HG011316, (L.C.), 1R01GM141627 (L.C.), R35DE030054 (J.B.S.), and R35GM155437 (R.A.H.-L.). L.C. is a Donald and Delia Baxter Foundation Faculty Scholar. R.A.H.-L.-L.holds a Career Award at the Scientific Interface from Burroughs Welcome Fund. R.A.H.-L.-L. is a Chan Zuckerberg Biohub San Francisco investigator, a Parker Institute for Cancer Immunotherapy Stanford Member researcher and a Pew Biomedical Scholar. R.K.D. was supported by the Stanford School of Medicine Dean’s Postdoctoral Fellowship and American Cancer Society Postdoctoral Fellowship.

## Supplementary Figures

**Fig S1:**
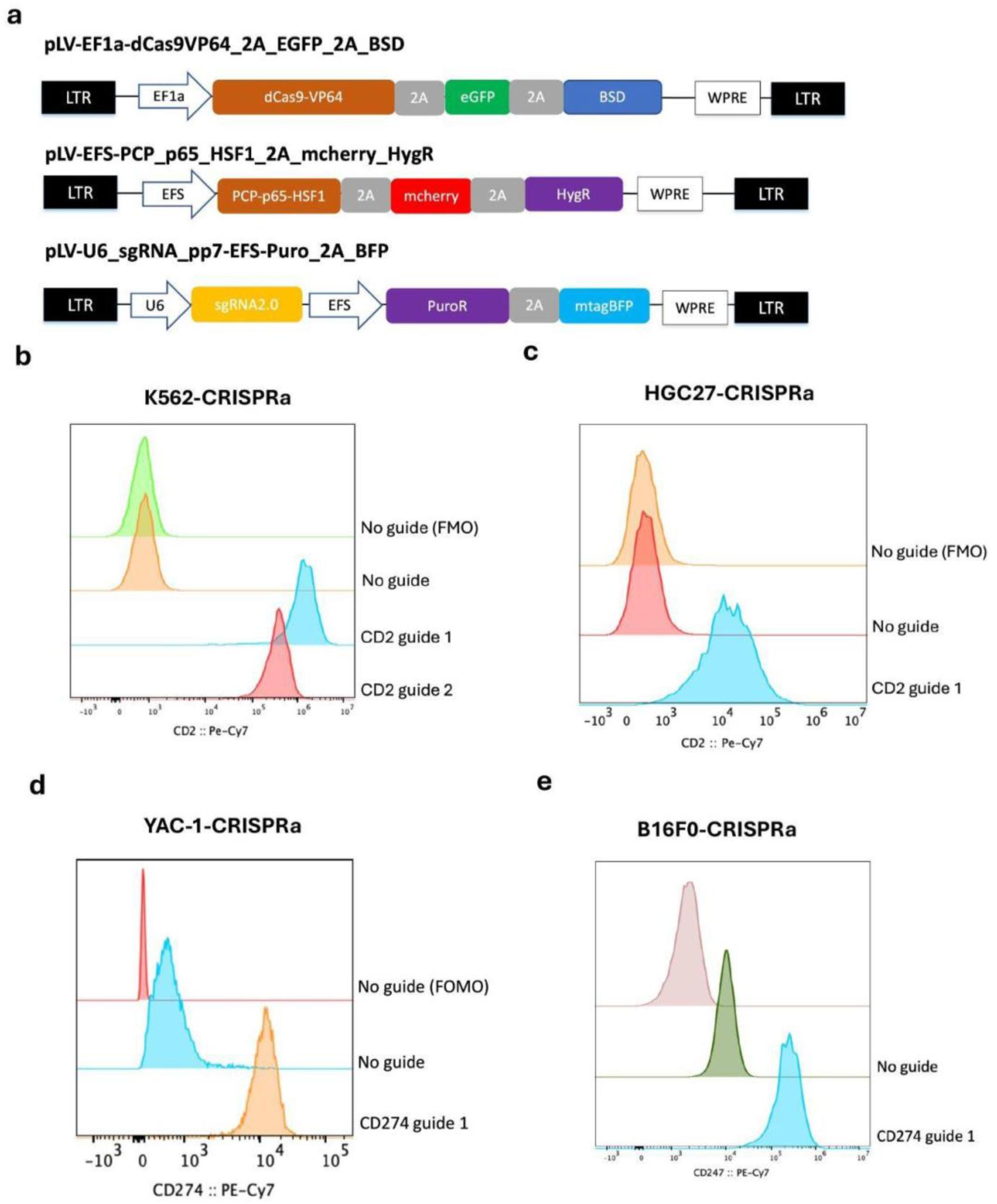
Design and validation of CRISPRa screening lines. **a,** Vectors for creation of CRISPRa activation cell lines. LTR– long terminal repeat; EF1a – Elongation factor 1 alpha promoter; 2A – 2A peptide; eGFP – enhanced green fluorescent protein; BSD – blasticdin deaminase ; WPRE – Woodchuck Hepatitis Virus posttranscriptional regulatory element; EFS – EF1a short promoter; PCP – PP7 coat protein; mcherry – monomeric cherry fluorescent protein; HygR - hygromycin resistance gene; U6 – human U6 promoter; PuroR – puromycin resistance gene; mtagBFP – monomeric tag blue fluorescent protein. **b-c,** Flow cytometry analysis of K562-CRISPRa **(b**) or HGC27- CRISPRa (**c**) transduced with a gRNA targeting the human *CD2* promoter. **d-e,** Flow cytometry analysis of YAC-1-CRISPRa (**d**) and B16F0-CRISPRa (**e**) cells transduced with a gRNA targeting the murine *Cd274* locus.

**Fig S2:**
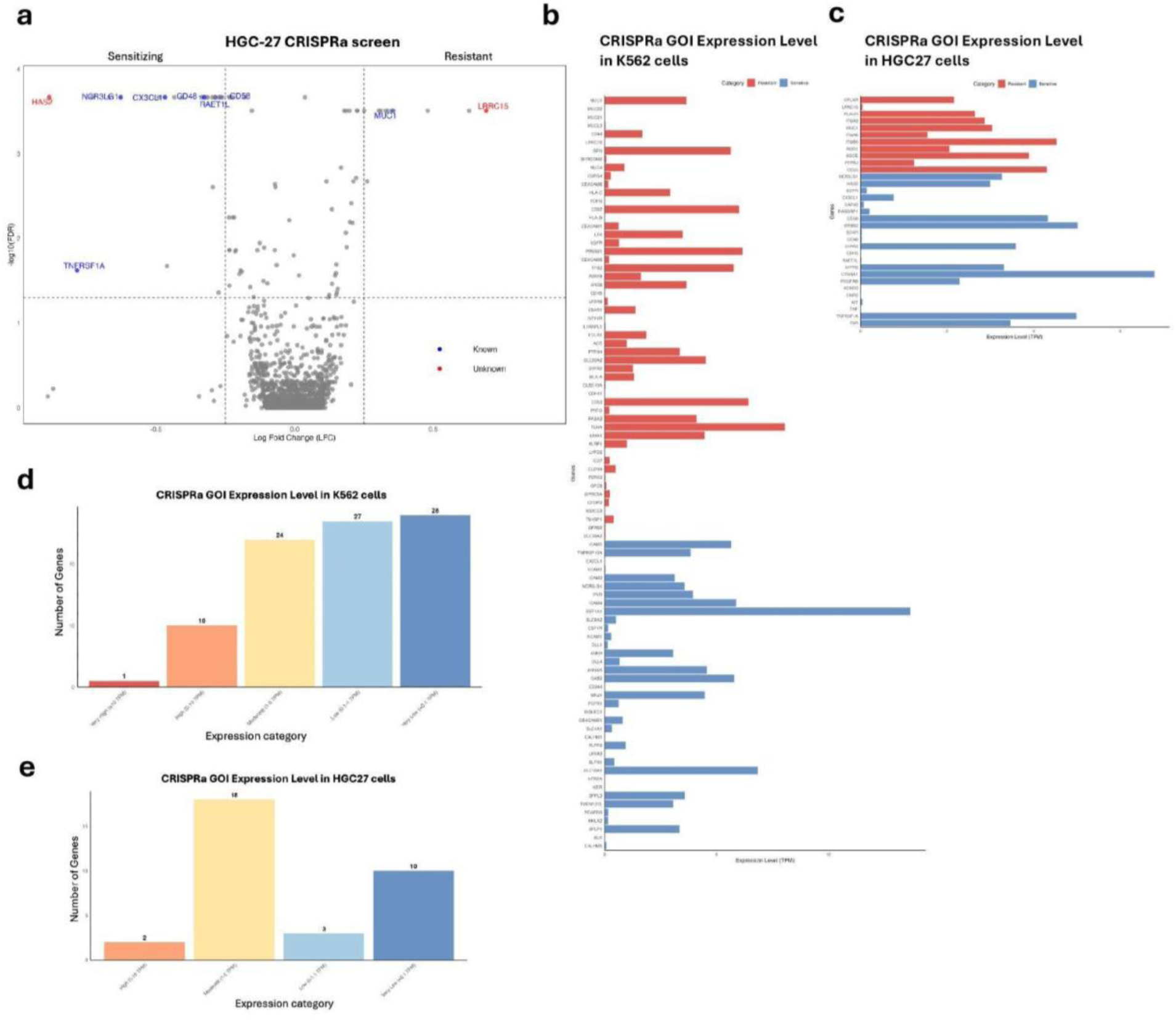
Analysis of human surfaceome CRISPRa screens. **a,** Volcano plot of surfaceome CRISPRa screen in HGC-27 cells. Known screen hits appear in blue and novel screen hits in red. **b-c,** Expression of candidate genes of interest from K562 (**b**) or HGC-27 (**c**) surfaceome CRISPRa screens within respective cell lines using DepMap RNA-Seq data. **d,e,** Candidate genes from K562 (**d**) or HGC-27 (**e**) categorized by levels of expression within respective cell lines using DepMap RNA-Seq data.

**Fig S3:**
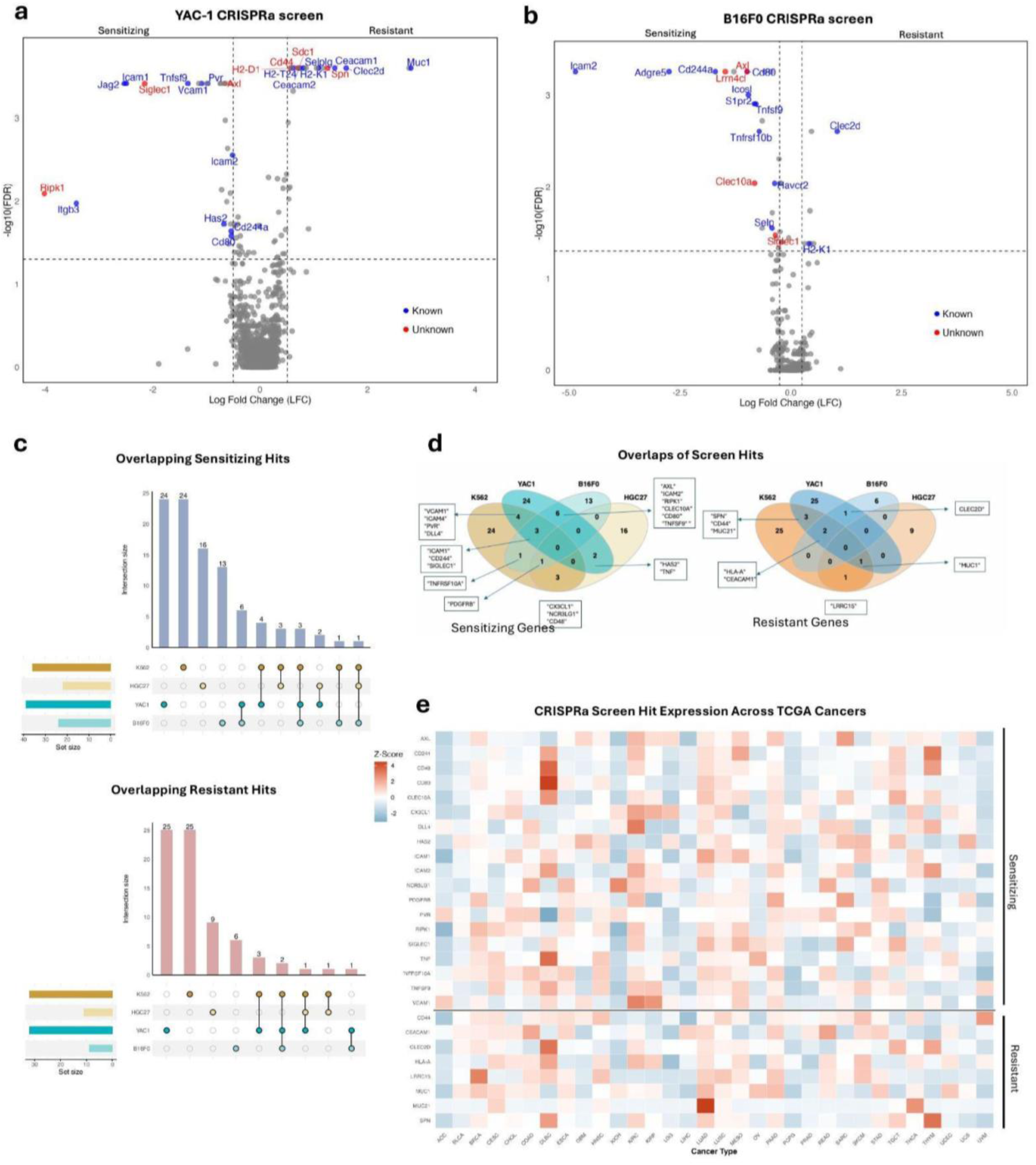
Analysis of mouse and human surfaceome CRISPRa screens. **a-b,** Volcano plots of surfaceome CRISPRa screens in YAC-1 (**a**) and B16F0 **(b**) cells. Known screen hits appear in blue and novel screen hits in red. **c,d** Overlapping sensitizing (**c**) and resistant (**d**) hits across four different surfaceome CRISPRa screens. **e,** Expression of common hits for CRISPRa screens across cancers in the TCGA database as z-scored log(TPM+1) data. Genes appear in at least 2 screens with the criteria of FDR < .05 and log2(FC) > 0.25. ACC - Adrenocortical carcinoma, BLCA -Bladder Urothelial Carcinoma, BRCA - Breast invasive carcinoma, CESC - Cervical squamous cell carcinoma and endocervical adenocarcinoma, CHOL - Cholangiocarcinoma, COAD - Colon adenocarcinoma, DLBC - Lymphoid Neoplasm Diffuse Large B-cell Lymphoma, ESCA - Esophageal carcinoma, GBM - Glioblastoma multiforme, HNSC - Head and Neck squamous cell carcinoma, KICH - Kidney Chromophobe, KIRC - Kidney renal clear cell carcinoma, KIRP - Kidney renal papillary cell carcinoma, LGG - Brain Lower Grade Glioma, LIHC - Liver hepatocellular carcinoma, LUAD - Lung adenocarcinoma, LUSC - Lung squamous cell carcinoma, MESO - Mesothelioma, OV - Ovarian serous cystadenocarcinoma, PAAD - Pancreatic adenocarcinoma, PCPG - Pheochromocytoma and Paraganglioma, PRAD - Prostate adenocarcinoma, READ - Rectum adenocarcinoma, SARC - Sarcoma, SKCM - Skin Cutaneous Melanoma, STAD - Stomach adenocarcinoma, TGCT - Testicular Germ Cell Tumors, THCA - Thyroid carcinoma, THYM - Thymoma, UCEC - Uterine Corpus Endometrial Carcinoma, UCS - Uterine Carcinosarcoma, UVM - Uveal Melanoma.

**Fig S4:**
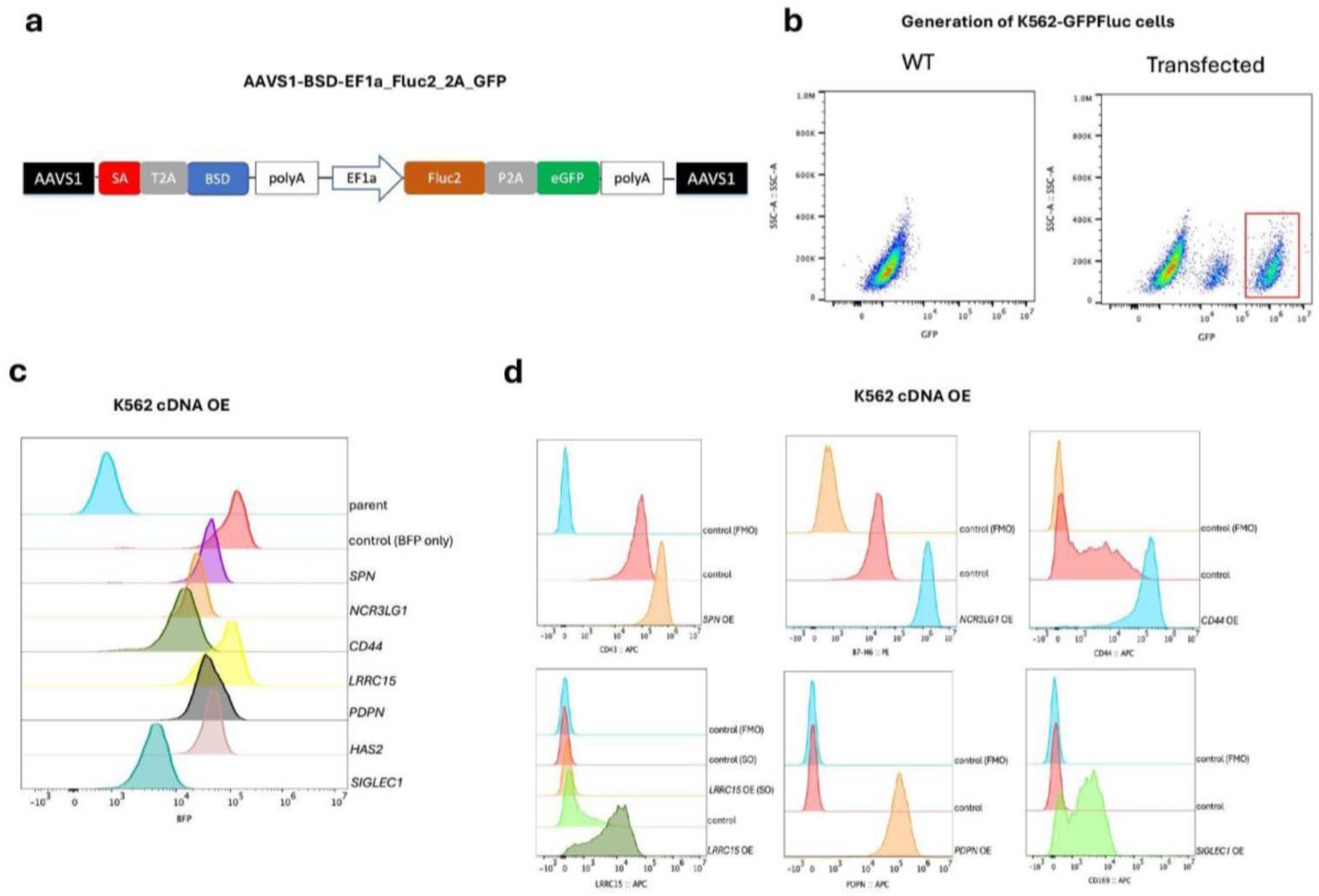
Cytotoxicity assays of cDNA overexpressing K562 cells. **a,** Design of FLuc2-2A-GFP AAVS1 knock-in (KI) construct. AAVS1 – 5’ and 3’ arms with genomic sequences from the human *AAVS1* locus; SA – splice acceptor site; 2A – 2A peptide; BSD – blasticidin deaminase; polyA – bovine growth hormone polyadenylation sequence; EF1a – Elongation factor 1 alpha promoter; Fluc2 – firefly luciferase 2; 2A – 2A peptide; eGFP – enhanced green fluorescent protein. **b,** Wildtype (WT) and FLuc2- 2A-GFP AAVS1 KI cells 14 days after transfection and selection. Cells gated in red were sorted and used for downstream assays. **c-d,** Flow cytometry of BFP (**c**) and cell surface ligand (**d**) expression in K562-GFPFluc cell transduced with cDNAs.

**Fig S5:**
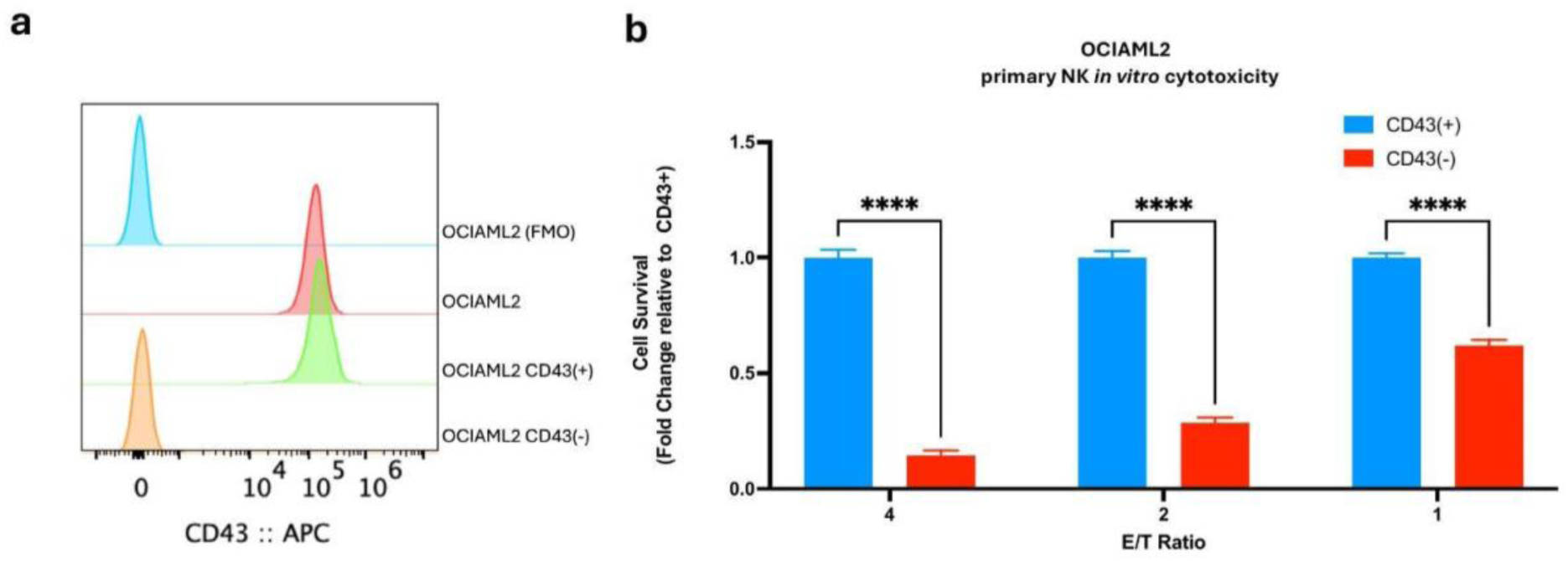
Effect of CD43 ablation on OCI-AML2 cells. **a,** Flow cytometry analysis plot of OCI-AML2 cells and derivatives for surface expression of CD43. **b,** NK cell cytotoxicity assays performed against CD43(+) and CD43(-) OCI-AML2 cells. Data normalized to CD43(+) for respective E/T ratio. n=2 independent experiments, 6 technical replicates each. Two-way ANOVA with correction for multiple comparisons. *, p<0.05; **, p<0.01; ***, p <0.001; ****, p<0.0001

**Fig S6:**
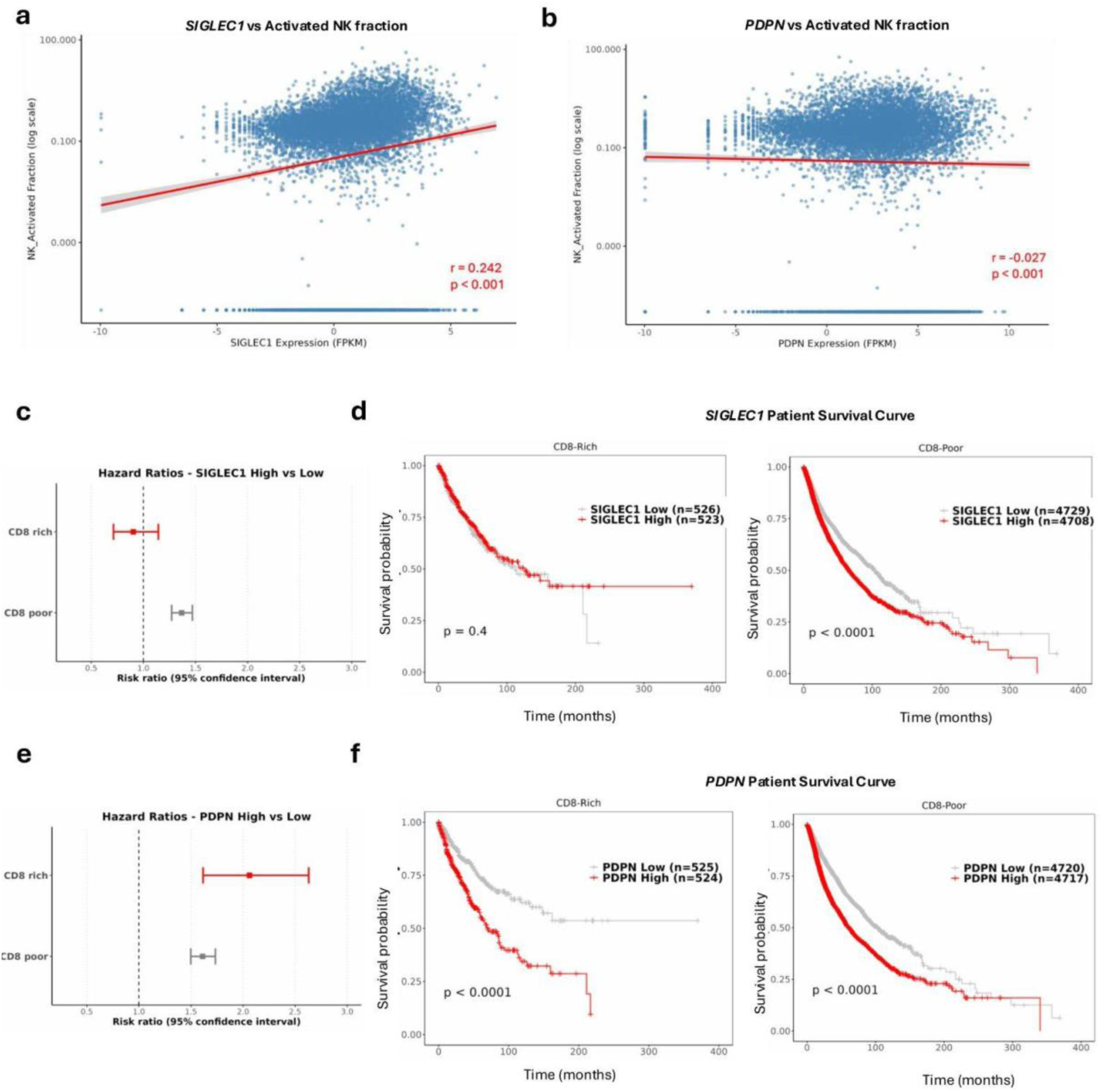
Correlates of gene expression on immune infiltration. **a,b,** Correlation between *SIGLEC1* (**a**) or *PDPN* (**b**) expression and NK activation state in TCGA cancers. n=10524. **c,e,** Cox hazard ratios of *SIGLEC1* (**c**) or *PDPN* (**e**) expression stratified by top and bottom 5% of CD8 T cell enrichment in TCGA cancers. **d,f,** Kaplan-Meier plot of *SIGLEC1* (**d**) or *PDPN* (**f**) expression stratified by top and bottom 5% of CD8 T cell enrichment in TCGA cancers.

**Fig S7:**
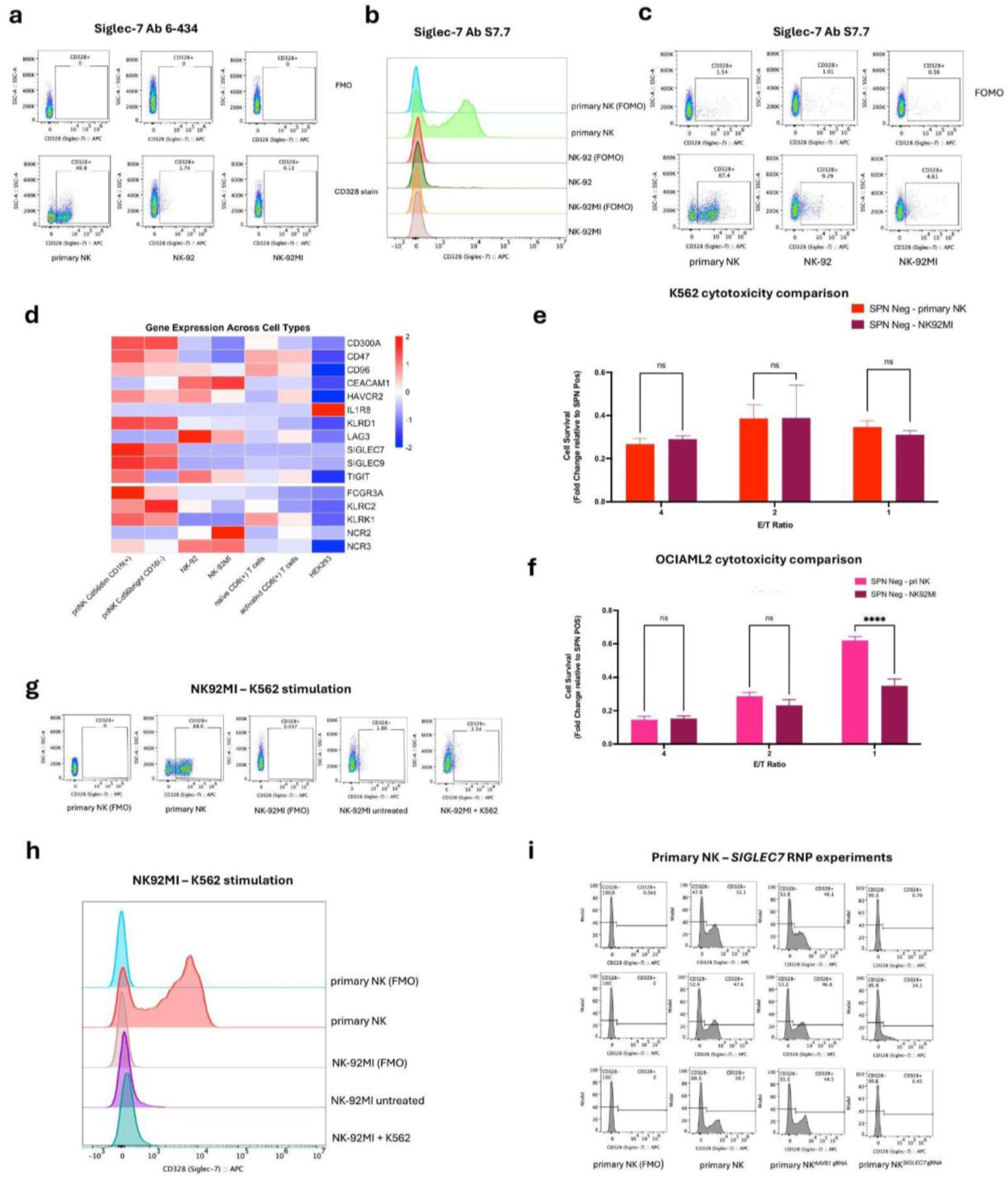
Role of Siglec-7 in CD43-mediated NK suppression. **a,** Flow cytometry analysis of CD328 expression using monoclonal Ab 6-434. **b-c,** Flow cytometry analysis of CD328 expression using monoclonal Ab S7.7. **d,** Heatmap of RNA-Seq expression of commonly expressed NK receptor genes. **e-f,** Comparison of relative fold differences in CD43(-)/CD43(+) for K562 (**e**) and OCI-AML2 (**f)** assayed against primary NK cells or NK-92MI. **g-h,** Representative flow cytometry analysis of CD328 expression on primary NK and NK-92MI cells co-cultured with K562 cells for 24 hr at 1:1 ratio. n=2 independent experiments. **i,** Flow cytometry plots of CD328 expression for all SIGLEC7/AAVS1 RNP targeting experiments. Cells assayed 4 days after nucleofection. **e-f,** n=3 independent experiments, six technical replicates. Two-way ANOVA with correction for multiple comparisons. *, p<0.05; **, p<0.01; ***, p <0.001; ****. ***.

**Fig S8:**
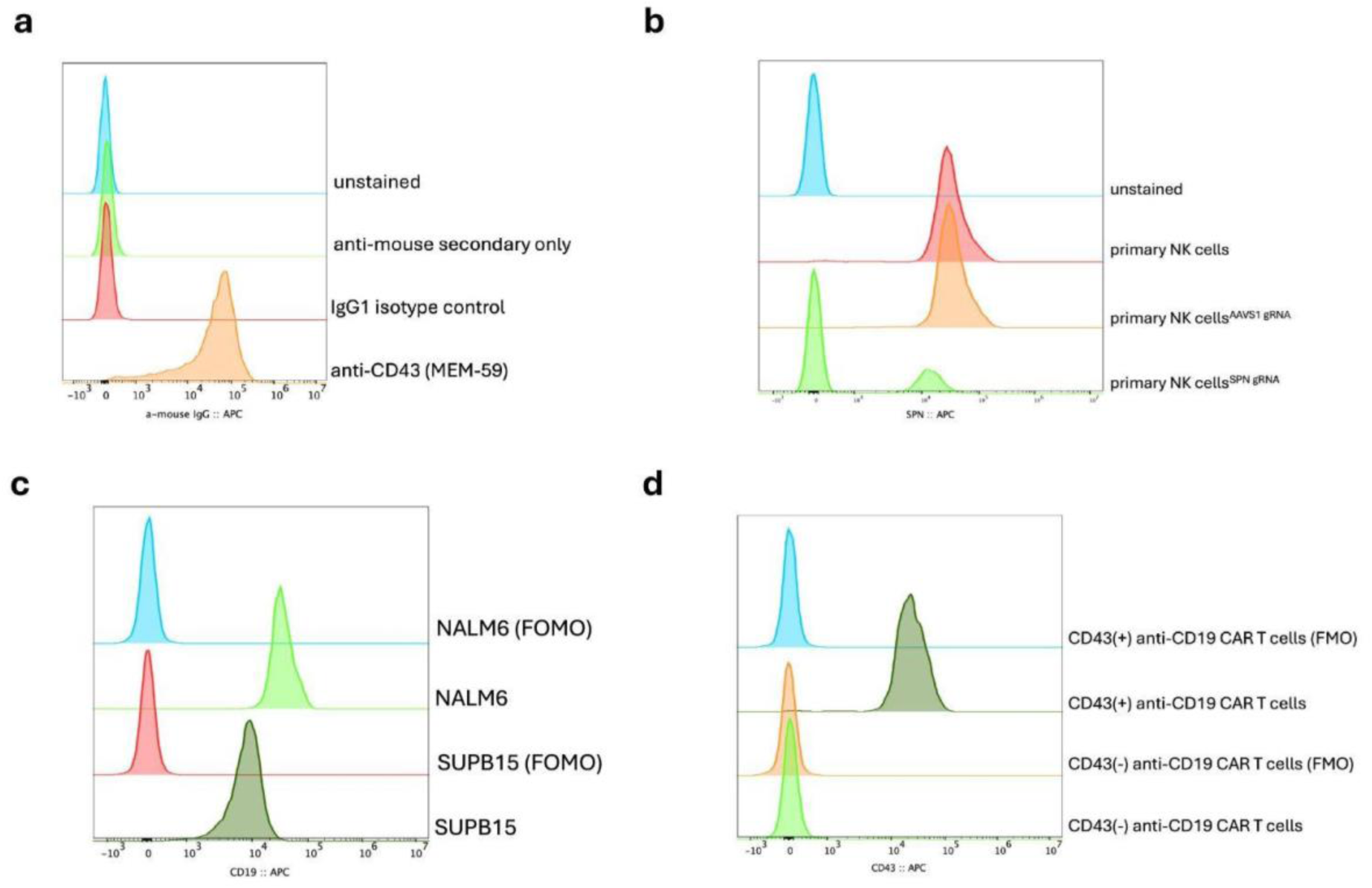
Targeting CD43 on NK cells and CAR T cells for improved tumor responses. **a,** Flow cytometry analysis of anti-CD43 antibody MEM-59 binding to primary NK cells. **b,** Flow cytometry analysis of CD43 expression in *AAVS1* and *SPN*-targeted primary NK cells. **c,** Flow cytometry analysis of CD19 expression on cancer lines. d, Flow cytometry analysis of CD43 expression on CD19-targeting CAR T cells electroporated with *AAVS1* and *SPN* RNP and sorted for CD43 expression.

